# N-glycosylation as a eukaryotic protective mechanism against protein aggregation

**DOI:** 10.1101/2023.08.11.552904

**Authors:** Ramon Duran-Romaña, Bert Houben, Matthias De Vleeschouwer, Nikolaos Louros, Matthew P Wilson, Gert Matthijs, Joost Schymkowitz, Frederic Rousseau

**Affiliations:** Switch Laboratory, VIB Center for Brain and Disease Research, 3000 Leuven, Belgium; Switch Laboratory, Department of Cellular and Molecular Medicine, KU Leuven, 3000 Leuven, Belgium; Laboratory for Molecular Diagnosis, Center for Human Genetics, KU Leuven, 3000 Leuven, Belgium

## Abstract

The tendency for proteins to form aggregates is an inherent part of every proteome and arises from the self-assembly of short protein segments called aggregation-prone regions (APRs). While post-translational modifications (PTMs) have been implicated in modulating protein aggregation, their direct role in APRs remains poorly understood. In this study, we used a combination of proteome-wide computational analyses and biochemical techniques to investigate the potential involvement of PTMs in aggregation regulation. Our findings reveal that while most PTM types are disfavored near APRs, N-glycosylation is enriched and evolutionarily selected, especially in proteins prone to misfolding. Experimentally, we show that N-glycosylation inhibits the aggregation of peptides *in vitro* through steric hindrance. Moreover, mining existing proteomics data, we find that the loss of N-glycans at the flanks of APRs leads to specific protein aggregation in Neuro2a cells. Our results point towards a novel intrinsic role for N-glycosylation, directly preventing protein aggregation in eukaryotes.

## INTRODUCTION

The conversion of soluble functional proteins into β-structured aggregates is triggered by short, generally hydrophobic, amino acid stretches known as aggregation-prone regions (APRs) [1]. Most proteins contain one and usually several APRs. In fact, around 20% of all residues in globular proteins are predicted to reside within these regions [2]. APRs are mostly buried inside the hydrophobic core of globular proteins, preventing them from initiating aggregation [3]. However, under physiological stress or during translation and translocation, APRs are exposed to the solvent and are prone to aggregate, requiring rigorous regulation by the cellular proteostasis machinery [4, 5]. Insoluble aggregates lead to the loss of function of the affected proteins and are often toxic to cells. This toxicity is strongly associated with a wide range of human diseases and ageing, including Alzheimer’s and Parkinson’s disease [6, 7].

The evolutionary persistence of APRs is a result of their necessity for protein stability, as the forces that drive aggregation, i.e., hydrophobicity and β-sheet propensity, are also crucial for the folding of globular proteins [8]. Nevertheless, throughout evolution, the potency of APRs has been minimised by the presence of adjacent residues that suppress aggregation propensity, known as aggregation gatekeepers [9]. Specifically, charged amino acids (Arg, Lys, Asp, and Glu) and proline (Pro) are significantly enriched in the regions immediately flanking APRs, as they kinetically and thermodynamically disfavour aggregation [2, 10–13]. The introduction of charges generates repulsion forces that strongly reduce aggregation propensity, while Pro is incompatible with the β-strand conformations associated with protein aggregation. Due to their anti-aggregation properties, gatekeepers are essential to maintain the overall fitness of cells, as they affect protein synthesis and degradation rates and can even act as molecular signals to recruit chaperones to non-native states [14, 15]. In fact, aggregation gatekeepers are evolutionarily conserved despite destabilising the native structure, showing that these residues constitute a functional class specifically devoted to proteostasis [16]. Accordingly, mutations that remove gatekeeper residues are more often associated with human diseases than neutral polymorphisms [17].

Many proteins are modified during or shortly after translation to assist protein folding and increase the stability of the native structure. Given this intimate connection with protein folding, it is perhaps unsurprising that protein post-translational modifications (PTMs) are gradually becoming associated with protein aggregation events. An increasing number of studies have shown that PTMs can directly – or indirectly – affect the aggregation potency of proteins associated with common aggregation diseases [18–20]. For example, phosphorylation interferes directly with Aβ fibrillary structure maturation [21], whereas in tau molecules, it reduces microtubule binding affinity, thus increasing the concentration of soluble tau and resulting in later-stage aggregation [22]. In recent years, the reversible O-GlcNAc modification has been shown to directly inhibit protein aggregation in many neurodegenerative diseases and indirectly promote cytoprotection against a wide range of cellular stresses [23, 24]. Nevertheless, it is unclear whether other PTM types constitute a general mechanism of aggregation prevention across proteomes.

The most abundant category of PTMs involves the enzymatic addition of functional groups to amino acid side chains, increasing their size and chemical complexity. Intriguingly, many PTM types have chemical properties reminiscent of gatekeeper residues as they often add bulk chains - likely incompatible with β-aggregation – and charges – potentially causing charge repulsion. In fact, negatively charged residues (Asp and Glu) have historically been used to mimic the phosphorylated state of proteins, as phosphorylation adds a negative charge to the amino acid side chain [25]. Furthermore, positively charged residues (Arg and Lys) are susceptible to many types of PTMs, such as acetylation or methylation. For these reasons, we hypothesise that PTMs could have been selected throughout evolution to intrinsically protect against aggregation, thus expanding the current repertoire of aggregation gatekeepers. In this work, we scanned the entire human proteome with a widely used protein aggregation prediction algorithm, TANGO, to analyse the frequency of the most abundant PTM types in APRs and their surrounding residues. Our findings show that N-glycosylation is significantly enriched, conserved, and commonly replaces unmodified gatekeeper residues at these positions. Using biophysical assays on N-glycosylated and unmodified aggregation-prone peptides, we show that this modification mitigates aggregation *in vitro* through steric hindrance. Analysis of the structural properties of proteins with APRs flanked by N-glycosylation indicates a preferential association with topologically complex domains that have a high aggregation propensity. Finally, re-analysis of proteomics data that measures changes in protein solubility after treatment of mouse Neuro2a cells with an N-glycosylation inhibitor shows the aggregation of specific newly synthesised proteins.

## RESULTS

### While most PTM types are disfavoured around APRs, N-glycosylation is enriched

Unmodified aggregation gatekeepers (Arg, Lys, Asp, Glu, and Pro) are significantly enriched in the positions immediately surrounding APRs. In fact, at least one of these amino acids is found within the three neighbouring residues – on either side – in more than 90% of all APRs identified by TANGO [1, 2], a widely used protein aggregation predictor. Therefore, to investigate the potential role of the most common types of PTMs as aggregation gatekeepers, we calculated their relative frequencies in and around human APRs. First, human proteins were scanned with TANGO, which identified 84,537 APRs (TANGO score >10 and length 5-15 residues). The three residues preceding and succeeding APRs were labelled as gatekeeping regions (GRs), and all other residues as distal regions (DRs) (**Figure 1A**). Next, experimentally annotated human PTM sites were collected from dbPTM [26] and the O-GlcNAcAtlas [27] and were mapped to the dataset. Only PTM types with sufficient data to perform accurate statistics were kept (at least 1,200 sites), which resulted in 17 PTM types across 571,759 unique sites (**Supplementary Table 1**).

**Figure 1.**
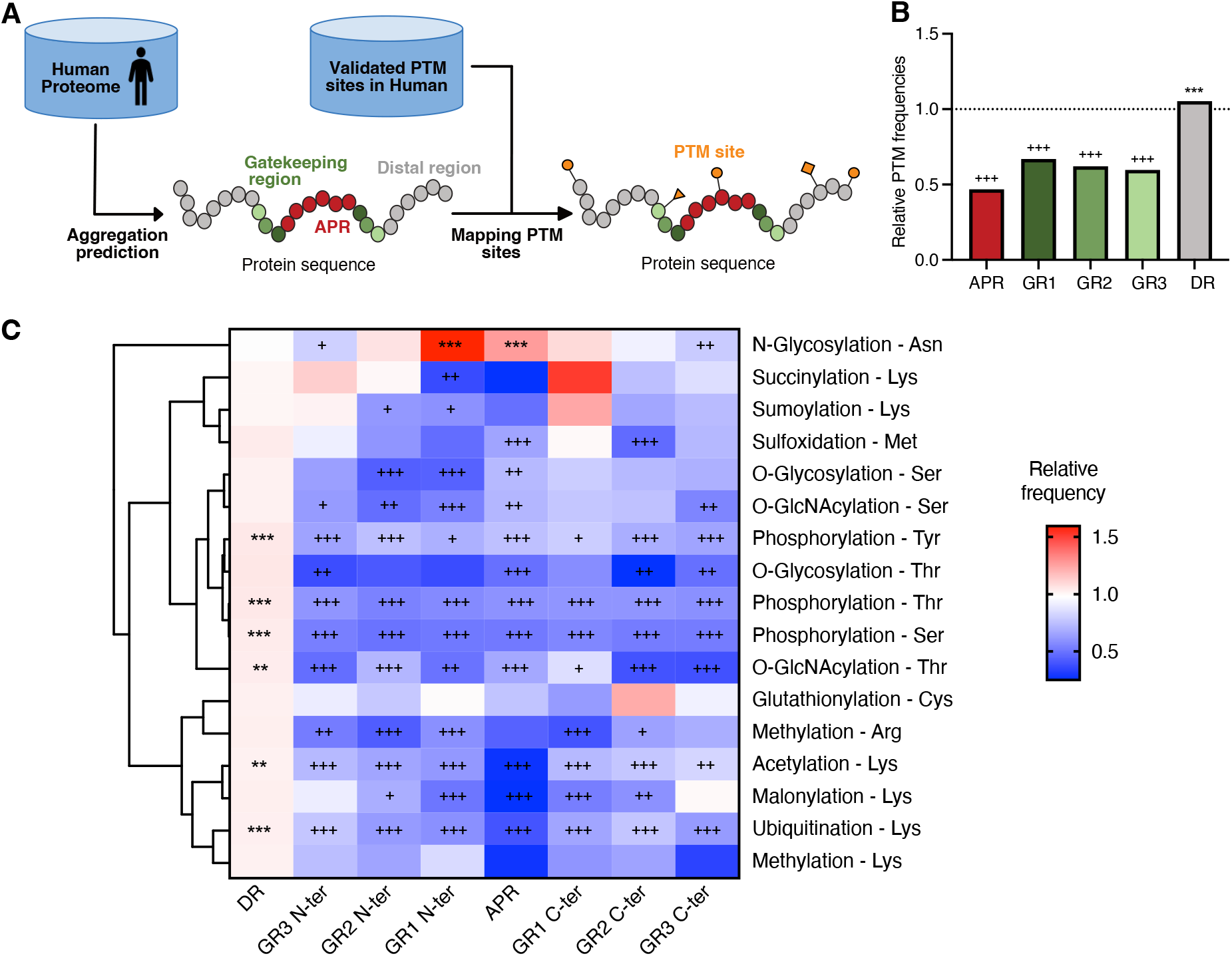
Relative enrichment of different PTM types in APRs and GRs. **A)** Schematic representation of the dataset preparation. **B)** Barplot showing the frequency of all PTM sites in APRs, GRs and DRs relative to background (all proteins containing the specific PTM). Crosses (asterisks) at the top of the bar indicate that a region has a significantly lower (higher) frequency compared to the background by Fisher exact test with FDR correction. **C)** Heatmap showing the relative frequencies for each of the 15 types of PTMs. Columns indicate the different protein regions, and rows show the PTM types. Statistics are calculated and illustrated as in B. Rows are clustered based on Pearson correlation as a distance measure.

Our findings show that PTMs, in general, are significantly underrepresented in APRs and GRs (**Figures 1B and 1C**), which means that most PTM types occur more frequently in residues that are located far away from APRs. This is not surprising as APRs are normally partially or completely buried in the folded structure, while PTM sites must be solvent accessible to be recognised by their modifying enzyme [28, 29] (**Supplementary Figure 1**). Nevertheless, restricting the analysis only to residues that are solvent accessible, and hence more readily modified, did not change these observations (**Supplementary Figure 2**). Another protein property that has been strongly associated with the occurrence of PTMs is structure disorder [29]. However, APRs and their GRs are predominantly found in structured domains, which could explain why PTM types that are more often observed in intrinsically disordered regions, such as phosphorylation or O-glycosylation, are disfavoured (**Supplementary Figures 3A and 3B**). This is also the case for O-GlcNAcylation, despite many reports showing that this modification dramatically slows down the aggregation of specific proteins involved in neurodegeneration, which are often highly disordered and polar [23]. Since disordered regions are depleted of a stable globular structure, their aggregation is driven more by β-sheet propensity rather than hydrophobicity [9].

In contrast to all other PTM types analysed, N-glycosylation is significantly enriched in APRs and GRs, especially at the N-terminal side (**Figure 1C**). Moreover, restraining the analysis only to exposed residues further increased this enrichment (**Supplementary Figure 2**).

### N-glycosites flanking APRs are evolutionarily conserved

N-glycosylation is one of the most common protein modifications in eukaryotic cells. It occurs in nearly all proteins that enter the secretory pathway (SP) and has essential roles in protein folding and quality control [30, 31]. The attachment of an N-glycan to an asparagine residue requires the recognition of a consensus sequence or sequon (Asn-X-Thr/Ser, where X ≠ Pro). This reaction is catalysed by an oligosaccharyltransferase (OST) on the luminal side of the endoplasmic reticulum (ER).

Since TANGO is a sequence-based predictor, we assessed whether the enrichment detected above was an artefact stemming from the Asn-X-Thr/Ser sequon being polar – and hence likely to be recognised as a gatekeeper when it flanks an APR – instead of a biological signal from the N-glycan. To check this, we compared the relative frequencies of sequons in proteins that have been experimentally determined to undergo N-glycosylation (SP glycosylated) to sequons that are either not glycosylated (SP non-glycosylated) or cannot be glycosylated due to their subcellular location (non-SP). An enrichment was only observed in APRs and GRs for glycosylated sequons (**Figure 2A**). This is highlighted in transmembrane proteins, as only those sequons in domains predicted to be in the extracellular or the lumenal side, which can therefore get glycosylated, showed an enrichment in these regions (**Supplementary Figures 4A and 4B**). Moreover, the enrichment was lost in sequons of artificial protein sequences that were randomly generated using the specific amino acid distributions of SP proteins (SP randomised), further indicating that it does not arise from sequence bias (**Figure 2A**). Finally, we observed a similar enrichment pattern when using a different aggregation predictor (CamSol [32]; **Supplementary Figure 4C**). Together, these results indicate that the enrichment of glycosylated sequons observed in APRs and GRs neither arises from a bias due to the sequon composition nor the choice of the aggregation predictor and, instead, is a direct result of N-glycosylation. Calculating the ratio between the relative frequencies of glycosylated sequons against the relative frequency of non-glycosylated sequons showed that there are three regions under positive selective pressure to be glycosylated, which we named enriched positions (EPs): GR2 N-ter, GR1 N-ter, and APR (**Figure 2B**). There are 1,155 N-glycosylated sites in EPs distributed in 858 unique proteins (15% of all SP proteins; **Figure 2C**). Analysis of the gene ontology terms of these proteins showed no overrepresentation of a particular biological function, suggesting that N-glycosylation in APRs and GRs is a general mechanism employed by a wide range of protein families (data not shown).

**Figure 2.**
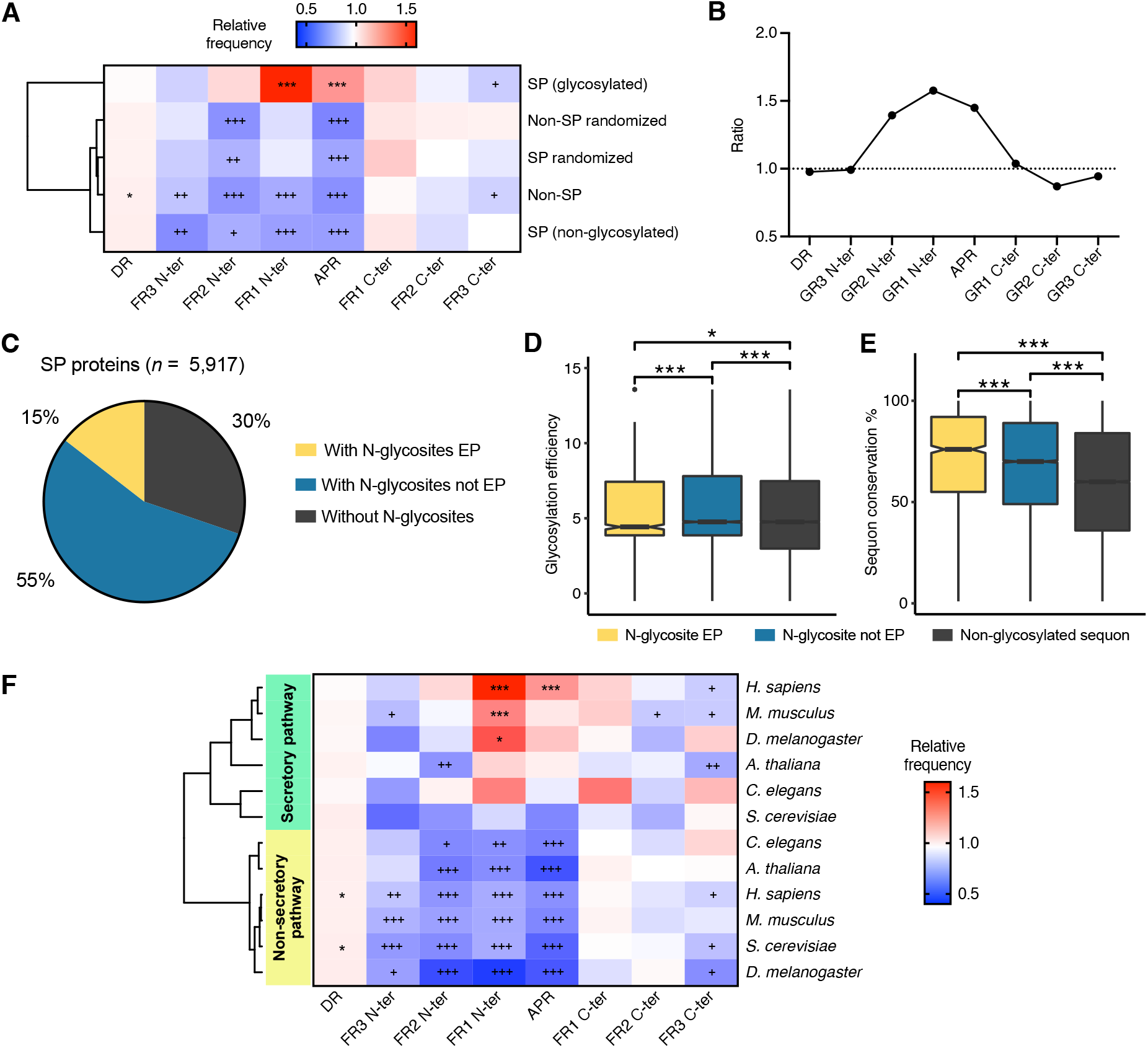
Functional assessment of N-glycosylation in APRs and GRs. **A)** Heatmap showing the relative frequencies of having a sequon in each region (columns) for different subsets of proteins (rows). Crosses (asterisks) at the top of the bar indicate that a region has a significantly lower (higher) frequency compared to the background by Fisher exact test with FDR correction. Rows are clustered based on Pearson correlation as a distance measure. **B)** Ratio between the relative frequencies of glycosylated sequons vs non-glycosylated sequons. **C)** Fraction of known secretory pathway proteins with at least one N-glycosylation site in enriched positions (yellow), with N-glycosylation sites that are not in enriched positions (blue) and without glycosylation sites (back). **D)** Boxplot showing the glycosylation efficiency of glycosylated sequons in enriched positions (yellow), rest of glycosylated sequons (blue) and non-glycosylated sequons (black) of human proteins. Unpaired Wilcoxon test was used to assess significance among groups with Bonferroni correction for multiple comparisons. **E)** Boxplot showing the conservation of human sequons in a set of 100 mammalian species for the same categories as D. Unpaired Wilcoxon test was used to assess significance among groups with Bonferroni correction for multiple comparisons. **F)** Heatmap showing the relative frequencies of having a sequon in each region for five different eukaryotic species. For all species, the relative frequencies of sequons in SP proteins are clustered together. The same is true for sequons in non-SP proteins. Statistics are calculated and illustrated as in A. Clustering is based on Pearson correlation as a distance measure.

N-glycosylation efficiency is highly influenced by the primary sequence context of glycosylation acceptor sites [33, 34]. Therefore, the specific sequence composition of APRs, GR2 N-ter and GR1 N-ter, could favour glycosylation efficiency. In other words, the strong selection observed at EPs might arise from the OST binding more efficiently to them instead of pointing to a shared functional role. To assess this, we predicted the glycosylation efficiency of human glycosylated sites using a model developed by Huang *et al*. [35]. In short, the authors used site-directed saturation mutagenesis to determine which residues improved or suppressed N-glycosylation efficiency. Based on their model, glycosylated sites in EPs have a lower efficiency compared to other glycosylated sites (**Figure 2D**), suggesting that the sequence composition of these regions is not driving their selection, and thus, hinting at an actual functional role. To corroborate this, we looked at the conservation of human sequons in a dataset of 100 mammalian species from the UCSC genome browser [36], as high conservation is commonly associated with an essential biological function. Indeed, N-glycosites in EPs have higher conservation compared to all other glycosylated sites, as well as to non-glycosylated sequons (**Figure 2E**).

The N-glycosylation pathway in the ER is very conserved across all eukaryotes [37, 38]. Therefore, we next investigated whether a similar enrichment pattern was present in other eukaryote model organisms. Given that the number of experimentally verified N-glycosites in other species besides human is very low, we assumed all sequons in SP proteins to be glycosylated. Strikingly, a similar enrichment pattern was found for sequons in SP proteins of other animals (*Mus musculus*, *Drosophila melanogaster,* and *Caenorhabditis elegans*) and plants (*Arabidopsis thaliana*), clustering together with the human SP enrichment profile (**Figure 2F**). Similarly, in these species, sequons of proteins that cannot get glycosylated (non-SP) were not enriched at EPs. For yeast (*Saccharomyces cerevisiae*), although its SP enrichment profile clustered together with the rest of the SP profiles, no enrichment was observed at these positions.

All of the above underlines a high selective pressure for N-glycosites in EPs to be preserved in evolution, pointing to a similar functional role for N-glycosylation in these sites. Since protein aggregation is generally detrimental for cells, we hypothesised that N-glycosylation is selected in these positions to protect against aggregation. In other words, this modification could be a novel class of aggregation gatekeeper.

### N-glycosites flanking APRs behave as and replace aggregation gatekeeper residues

The presence and number of unmodified gatekeeping residues (Arg, Lys, Asp, Glu, and Pro) flanking an APR correlate strongly with its aggregation propensity [39]. To investigate whether N-glycosites flanking APRs act as aggregation gatekeepers, we analysed the aggregation propensity (TANGO score) of APRs containing glycosylated and non-glycosylated sequons at EPs. We see that APRs flanked by N-terminally glycosylated sequons at GR1 N-ter and GR2 N-ter have significantly higher aggregation propensities than those flanked by non-glycosylated sequons in the same positions (**Figure 3A**). However, despite having a higher aggregation propensity on average, these APRs are flanked by fewer unmodified gatekeeping residues (**Figure 3B**). In fact, while in non-glycosylated sequons the number of unmodified gatekeepers increases with APR strength, in glycosylated sequons, the number remains low and constant across different APR strength bins (Supplementary Figure 5A). Since unmodified gatekeeping residues are crucial to avoid aggregation, especially for very strong APRs, this data suggests that N-glycans are replacing them in these positions, thus potentially taking their function as aggregation breakers. In contrast, none of the other GRs showed a significant difference in aggregation propensity or in the number of flanking unmodified gatekeeping residues (**Supplementary Figures 5B and 5C**). Glycosylated sequons in APRs did not show a difference in any of these analyses either (**Figures 3A and 3B**), despite being under positive selective pressure. A possible explanation is that APRs comprise a much larger region (5-15 aa), which adds noise to the analysis.

**Figure 3.**
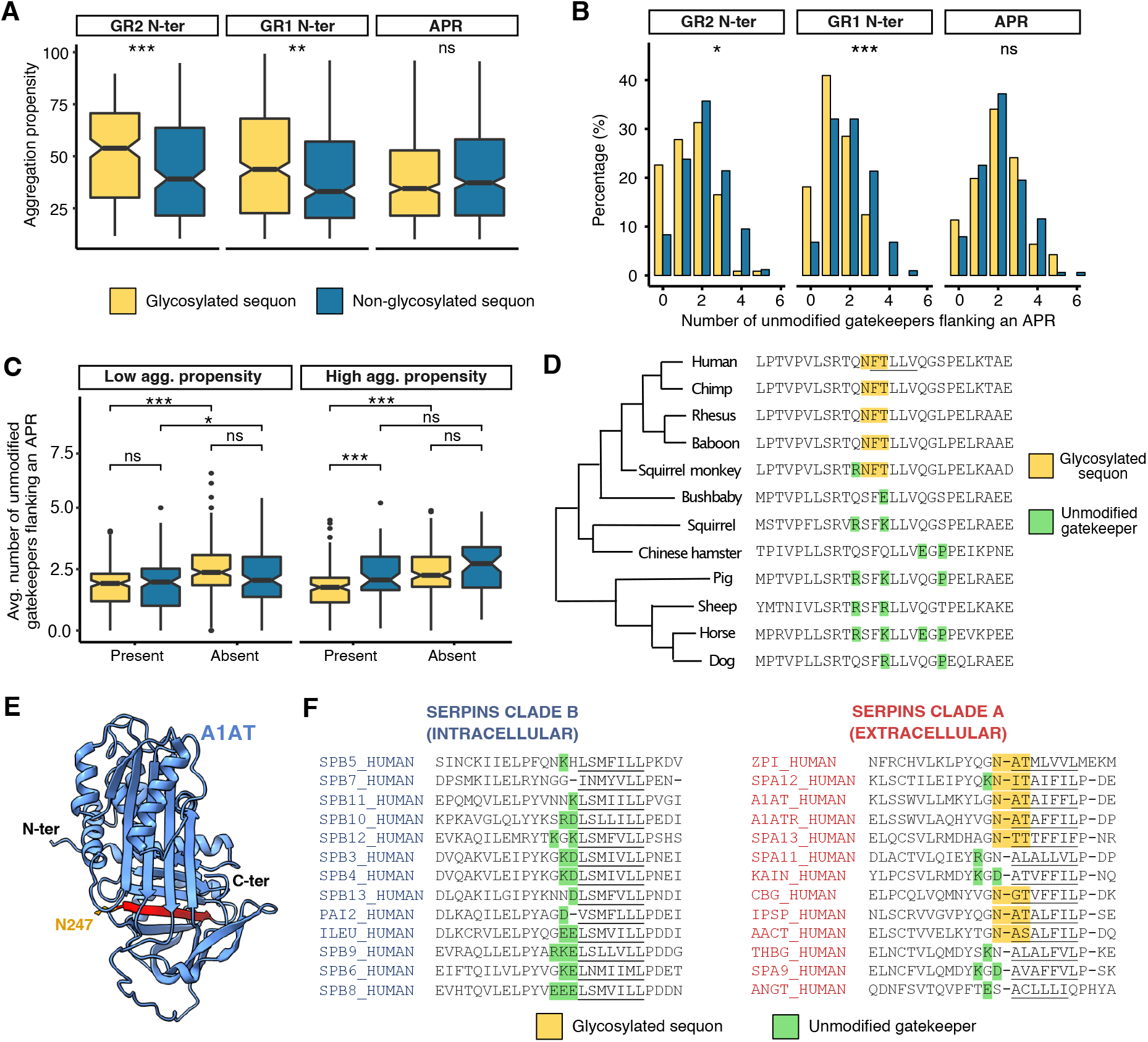
N-glycosites at EPs behave as aggregation gatekeepers. **A)** Boxplot showing the aggregation propensity (TANGO scores) of APRs that have glycosylated sequons or non-glycosylated sequons for each of the three EPs (GR2 N-ter, GR1 N-ter and APR). Unpaired Wilcoxon test was used to assess significance between the two groups (glycosylated vs non-glycosylated sequons). **B)** Barplot indicating the distribution of the number of charged residues in the three positions upstream and downstream of APRs with an aggregation propensity score >= 50. This threshold was used to ensure a strong evolutionary pressure to mitigate the aggregation of the APRs. Unpaired Wilcoxon test was used to assess significance between the two groups. **C)** Boxplot showing the average number of unmodified gatekeepers flanking glycosylated and non-glycosylated sequons in GR1 N-ter when these are present or absent throughout mammalian evolution. APRs are divided into two categories: weak if the TANGO score is < 50 or strong if the TANGO score is >= 50. **D)** Small subset of the multiple sequence alignment for BCAM. Glycosylated sequons are highlighted in yellow and unmodified gatekeepers are highlighted in green. The human APR is underscored. **E)** Example of a serpin structure (Alpha-1 antitrypsin; A1AT) obtained from AlphaFold and with the conserved aggregation-prone region highlighted in red. This particular serpin has an N-glycosylated site flanking the APR (orange). **F)** Multiple sequence alignment showing the same region for intracellular (clade B) and extracellular (clade A) serpins. The conserved APR is underscored in each protein. N-glycosylated sites or unmodified gatekeepers three residues upstream of the APR are highlighted in yellow or green, respectively.

To gain more insight into the role of N-glycosites as gatekeepers of aggregation, we looked at the conservation of human glycosylated and non-glycosylated sequons throughout mammalian evolution. Particularly, we focused on sequons at GR1 N-ter since this is the position that showed the highest enrichment and strongest selective pressure when it is glycosylated (**Figures 2A and 2B**). Each sequon at GR1 N-ter was mapped to the multiz100way dataset [36], a dataset containing multiple sequence alignments of 100 mammalian species to the human genome. We then calculated separately the average number of unmodified gatekeepers in protein orthologs for which the sequon is present and orthologs for which it is absent. In agreement with our previous analysis, we found that when glycosylated sequons acting as gatekeepers are present in a species, these are usually flanked by only a small number of unmodified gatekeepers, even when placed next to very strong APRs (**Figure 3C**). However, a significantly higher number of unmodified gatekeepers are found flanking APRs when glycosylated sequons are not present in a species. An example of this can be seen in the basal cell adhesion molecule protein (BCAM; **Figure 3D**). Non-glycosylated sequons are already flanked by a higher number of unmodified gatekeepers, particularly in the case of strong ARPs and, therefore, their absence in a species does not lead to an increase of unmodified gatekeepers (**Figure 3C**).

A similar observation was obtained when analysing protein paralogs, particularly the serpin superfamily of protease inhibitors. In humans, most serpins are classified into two clades: the extracellular ‘clade A’ and the intracellular ‘clade B’ [40, 41]. Interestingly, we found that many extracellular serpins have a glycosylated sequon flanking a very strong APR that is conserved in both clades (**Figures 3E and 3F**). However, in intracellular serpins, this APR is flanked instead by one or more unmodified gatekeeping residues, evidencing again an analogous function for N-glycans and unmodified gatekeepers (**Figure 3F**).

### N-glycosylation efficiently inhibits peptide aggregation *in vitro* by steric hindrance

The bioinformatics analysis presented above hints at a protective role of N-glycosylation against the aggregation of its cognate APRs. To experimentally assess this, we measured the aggregation kinetics and solubility of peptides with and without an N-glycan (**Figure 4A**). Short aggregating peptides were used instead of full proteins to mimic exposed APRs and to ensure the interpretability of our findings.

**Figure 4.**
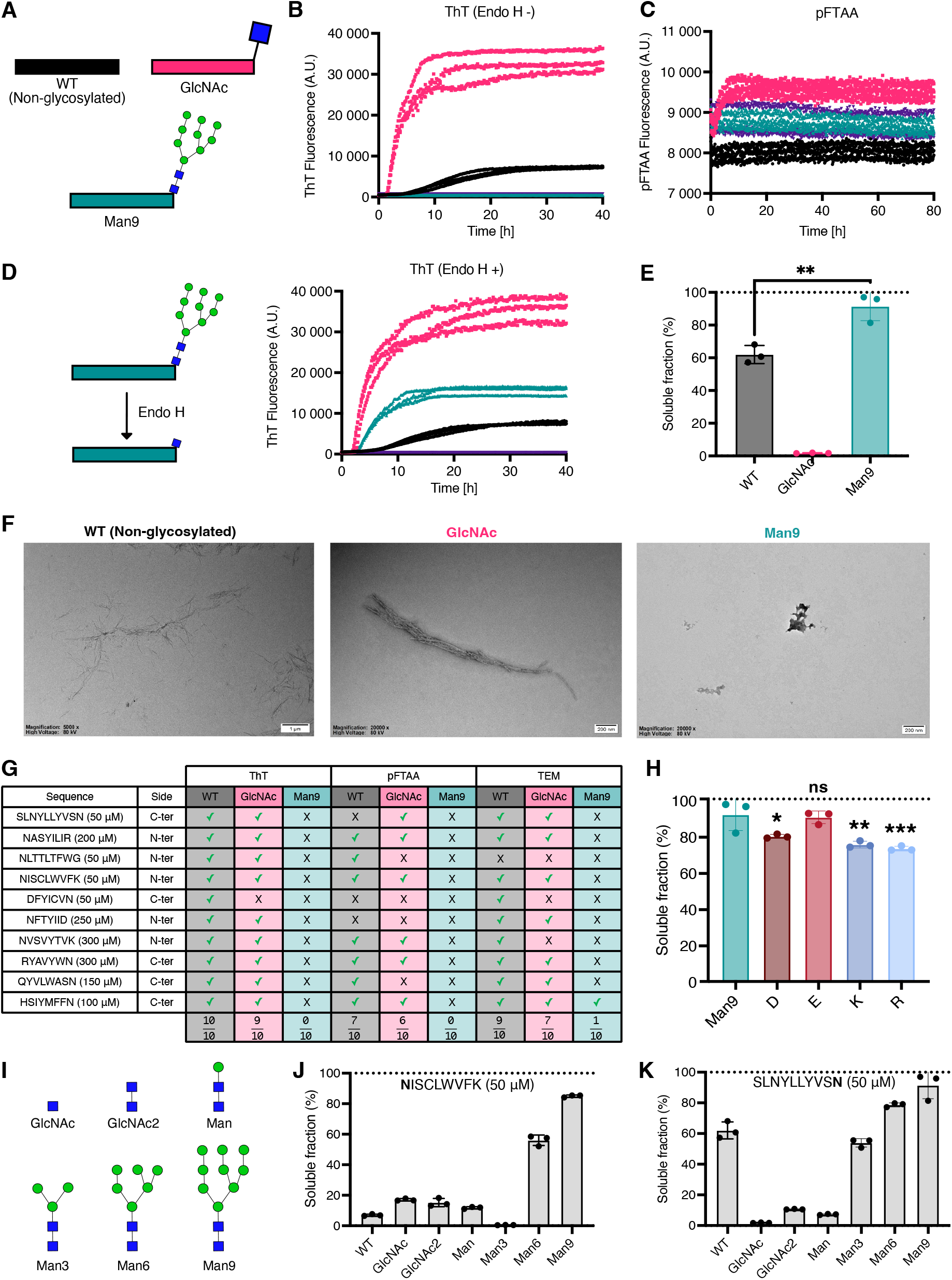
In vitro analysis of N-glycosylated peptides. **A)** Schematic representation of the peptide variants and experimental design. An aggregation core is flanked by either a non-glycosylated Asn (WT), GlcNAc or Man_9_. **B,C)** ThT binding (B) and pFTAA binding (C) kinetics of the SLNYLLYVSN peptide set. Fluorescence over time is shown for three independent repeats. Vehicle control fluorescence is shown in purple. **D)** ThT binding after incubation with 1 μL (500 units) of Endo H enzyme, which cleaves the bond between two N-acetylglucosamine (GlcNAc) subunits directly proximal to the asparagine residue of the glycopeptide. Fluorescence over time is shown for three independent repeats. Vehicle control fluorescence is shown in purple. **E)** Percentage of the concentration of peptide in the soluble fraction after ultracentrifugation for the SLNYLLYVSN peptide set (n=3). Unpaired t-test was used to assess significance. **F)** TEM images for the SLNYLLYVSN peptide set after seven days of incubation. **G)** Combined results for all APRs. Peptides were classified on whether they showed kinetics (ThT and pFTAA) and whether they formed fibrillar aggregates detectable by TEM imaging. **H)** Percentage of soluble fraction for the charged residue variants (D, E, K and R; n= 3). Man_9_ values were re-used from E. Unpaired t-test was used to assess significance against Man9. **I)** Schematic representation of the structures of the different glycoforms analysed. **J,K)** Percentage of soluble fraction after ultracentrifugation for the non-glycosylated and glycoforms versions of NISCLWVFK (J) and SLNYLLYVSN (K) peptide sets (n= 3). Non-glycosylated and Man_9_ peptides values were re-used from E and Supplementary Figure 13.

After an N-glycan precursor (Glc_3_Man_9_GlcNAc_2_) is transferred to a protein, it is processed in the ER by removal of the glucose residues as part of the quality-control process [30]. Then, the protein moves to the Golgi apparatus, where the carbohydrate is further processed into an extensive array of mature and complex N-glycoforms [42]. This raises the question whether there is a particular glycoform that confers protection against aggregation or, instead, if it is an intrinsic effect of all glycoforms. The genomes of higher eukaryotes encode two STT3 proteins (STT3A and STT3B), which are the catalytic subunits of two distinct OST complexes [37]. The STT3A complex is associated with the protein translocation channel and glycosylates the majority of sites as they emerge into the ER lumen while specific glycosites that are skipped by the STT3A complex are modified post-translationally by the STT3B complex. In other words, the addition of most N-glycans takes place while a protein is being translated and, therefore, before it folds [43]. During this time, an APR is exposed and at risk of engaging in non-native interactions, such as aggregation. Therefore, we reasoned that this is the most vulnerable time point in a protein lifespan – when it is most in need of anti-aggregation mechanisms – and decided to use the Man_9_GlcNAc_2_ (Man_9_) glycoform since it is the minimal carbohydrate structure that is attached to the nascent polypeptide before its folding.

We analysed ten human APRs with a flanking N-glycosite (**Supplementary Table 2**). In order to investigate if any structural constraints explain why the enrichment in our previous analysis was only observed in the N-terminal flank, we chose five APRs that were modified in the N-terminal site and five in the C-terminal site. Man_9_ variants for each APR were compared to their unmodified versions. In addition, GlcNAc versions of each peptide were made to determine if shorter N-glycan forms can inhibit aggregation. All peptides in a set were dissolved to a concentration in which the unmodified variant displayed dye-binding aggregation kinetics with Thioflavin-T (ThT). The results for the peptide set derived from SLNYLLYVSN are shown in **Figures 4B-4H**. ThT- and PFTAA-binding experiments revealed that aggregates were formed by the non-glycosylated and GlcNAc peptides, while for Man_9,_ no fluorescent signal was observed (**Figure 4B and 4C**). Incubating the Man_9_ peptide with Endo H, an enzyme that catalyses the conversion of Man_9_ into GlcNAc, resulted in a strong ThT fluorescent signal (**Figure 4D**), suggesting that the Man_9_ glycoform was inhibiting aggregation. However, since Man_9_ is a huge molecule, its size could hinder the binding of the fluorescent dyes to a potential aggregated structure. In order to dismiss this possibility, we used an orthogonal assay that measures the concentration of soluble peptide left once the aggregation reaction has reached an equilibrium. In short, peptides were incubated for a week and then subjected to ultracentrifugation. Endpoint solubility measurements of this peptide set showed that Man_9_ substantially improved APR solubility compared to non-glycosylated and GlcNAc peptides (**Figure 4E**). We reached similar conclusions by TEM imaging where no aggregated species were observed for the Man_9_ peptide, while both non-glycosylated and GlcNAc peptides formed amyloid fibrillar structures (**Figure 4F**). Together, these results indicate that Man_9_ strongly inhibits the formation of aggregates. The combined results of the ten APRs analysed confirmed the generality of these findings (**Figure 4G and Supplementary Figures 6-14**). Next, we made peptides in which the modified Asn residue was replaced by each of the four charged residues (D, E, R, and K) since these are known to strongly oppose aggregation. For the SLNYLLYVSN peptide set, Man_9_ was more soluble than all peptide versions with charged residues, apart from Glu (**Figure 4H**). Furthermore, in each APR set, Man_9_ was as good or better at improving the solubility of peptides compared to their charged counterparts (**Supplementary Figures 6-14**). This enhanced solubility could partially explain why N-glycosylation is selected over unmodified gatekeeping residues in some proteins. Surprisingly, while the computational analysis showed selection only for N-glycosites at the N-terminal flanks of APRs, the *in vitro* experiments revealed that N-glycosylation can inhibit aggregation in both flanks. This indicates that the preference for N-terminal flanks observed computationally does not arise from any APR-intrinsic structural constraints and, therefore, other biological factors may be responsible (see **Discussion**).

While Man_9_ showed complete or strong inhibition of aggregation in all peptide sets, GlcNAc’s capability of inhibiting aggregation was significantly lower. Moreover, in some peptide sets, GlcNAc actually enhanced aggregation (**Figure 4 and Supplementary Figure 13**). Previous studies have proposed that the large size and hydrophilicity of glycans prevent the aggregation of protein pharmaceutical products through steric hindrance [44, 45]. Therefore, we hypothesised that the difference in size between the two glycoforms might be responsible for the degree of inhibition observed. To assess this, we measured, in two of the peptide sets, the solubility of four additional glycoforms: GlcNAc_2_, ManGlcNAc_2_ (Man), Man_3_GlcNAc_2_ (Man_3_), and Man_6_GlcNAc_2_ (Man_6_) (**Figure 4I**). Interestingly, GlcNAc, GlcNAc_2_ and Man caused a minor and similar increase in solubility for the NISCLWVFK peptide compared to its unmodified version (**Figure 4J**), and were actually found to be more insoluble for the SLNYLLYVSN peptide (**Figure 4K**). A possible explanation could be the presence glycoform-specific interactions, leading to stacking between the hydrophobic faces of sugars or between aromatic residues and sugars of different peptides [46]. Conversely, Man_6_ and Man_9_ caused a substantial and size-dependent increase in solubility in both peptide sets (**Figure 4J and Figure 4K**), supporting that steric hindrance may be the mechanism behind aggregation inhibition. These results provide direct evidence that different glycoforms confer distinct levels of protection against aggregation. Moreover, the more potent inhibitory effect of Man_9_ on aggregation supports the idea that N-glycan-mediated protection against aggregation occurs before protein folding in the ER.

### N-glycosylation protects against aggregation in hard to fold proteins

Out of all APRs in proteins that follow the SP, only around 7% are flanked by N-glycans at EPs (**Figure 5A**). Why do some APRs, or the proteins bearing those APRs, require the extra level of protection granted by N-glycosylation? To answer this, we built a random forest classifier that predicts which APRs are protected by N-glycans using different features related to structural topology and aggregation, both at the APR and protein domain levels (see **Methods**). We decided to use features of individual protein domains instead of features from full proteins, as domains are independent evolutionary units that often fold independently from each other [47]. Domains were extracted using CATH-Gene3D [48, 49]. Since the number of protected and unprotected APRs is quite different and random forests are known to be sensitive to class imbalance, we trained two different models with opposite resampling techniques. Interestingly, the relative contact order of a domain was the most important feature in both models (**Figure 5B** and Supplementary Figure 15A). This is a widely used metric to describe the complexity of a polypeptide fold, which correlates with folding times [50]. Indeed, when comparing domains with at least one APR, those with an N-glycosite at EPs have a significantly higher relative contact order (**Supplementary Figure 15B**). Moreover, while high contact order domains without protected APRs generally have lower aggregation propensities, the ones with N-glycosites at EPs usually contain much stronger APRs (**Figure 5C**). Thus, N-glycosylation is placed in APRs of complex domains with overall high aggregation propensities. As expected, other parameters determined to be important by both models were the solvent accessibility of APRs and the number of unmodified gatekeeping residues flanking them (**Figure 5B and Supplementary Figures 15C and 15D**). N-glycosylation constrains part of the APR to be solvent accessible to avoid steric clashes, while from our previous analyses, we know that N-glycosylation replaces unmodified gatekeeping residues at EPs. The oxidising environment of the ER allows for the formation of disulphide bridges, which help stabilise the native fold of SP proteins. Nevertheless, the number of disulphide bridges in a domain had low importance in the prediction (**Figure 5B**).

**Figure 5.**
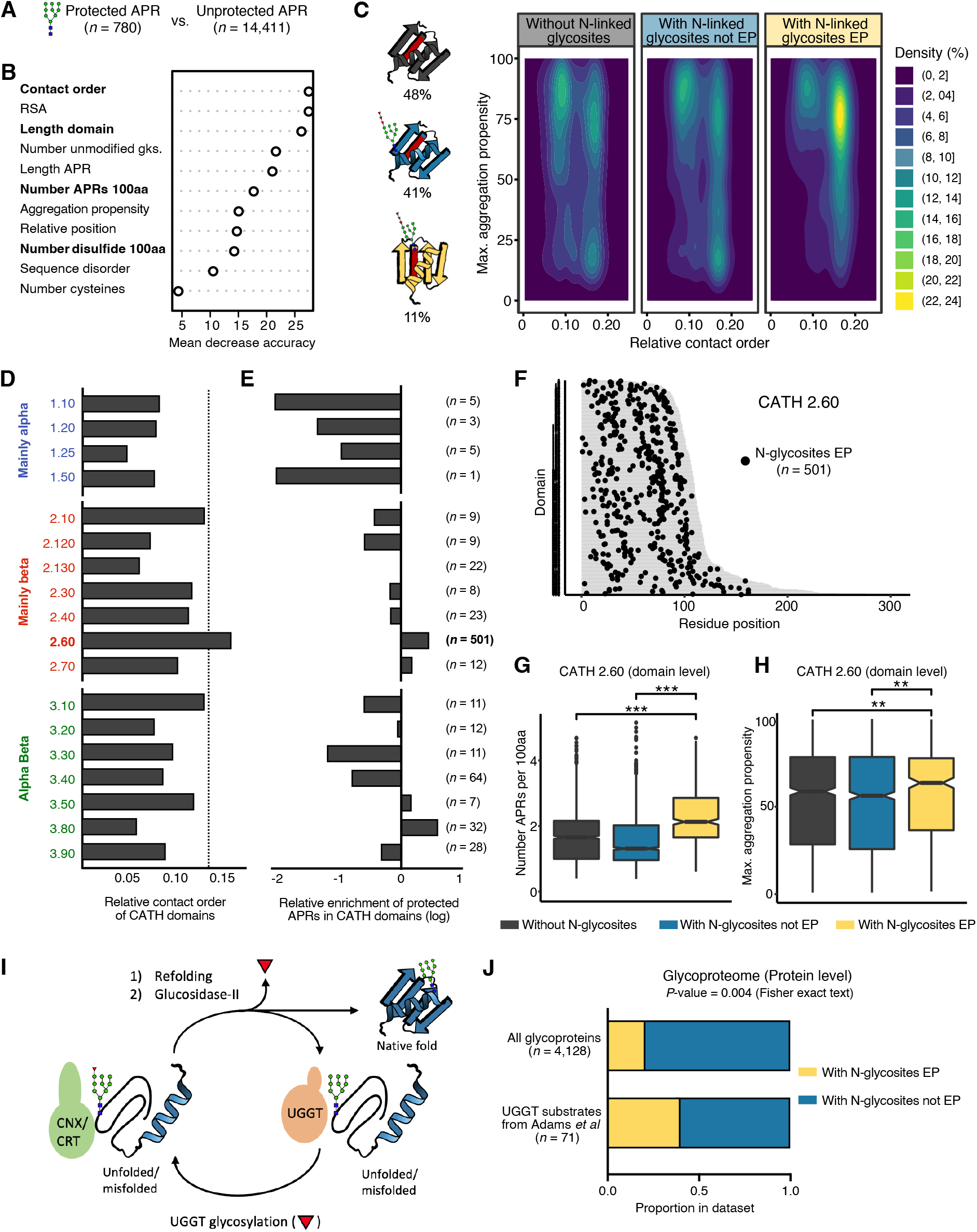
N-glycosylation protects against aggregation in hard-to-fold proteins. **A)** A random forest classifier was built to classify APRs present in CATH domains as protected (with N-glycosylation sites in EPs) and unprotected (all others). **B)** Variable importance plot for the model built using random undersampling. Mean accuracy indicates the performance of the model after removing a specific variable. Higher values indicate more importance of that variable in predicting protected vs unprotected APRs. Domain-specific variables are highlighted in bold, while APR-specific variables are unhighlighted. **C)** On the right, a two-dimensional density plot showing the relative contact order and the maximum aggregation propensity for domains classified in three categories: with N-glycosites in EPs (yellow), with N-glycosites not EPs (blue) and without N-glycosylated sites (black). On the left, a schematic representation of each domain category together with their percentage in the dataset. **D)** Average relative contact order of domains in each CATH architecture. The dotted line indicates the average relative contact order of domains containing an N-glycosite in an EP. **E)** Relative frequencies of finding a protected APR in each CATH architecture. The number of protected APRs present in each CATH architecture is shown. **F)** Map showing the position of N-glycosylation sites at EPs in all β-sandwich domains. Domains are sorted by length and coloured in grey. **G)** Boxplot showing the number of APRs per 100 amino acids in β-sandwich domains with N-glycosites in EPs (yellow), with N-glycosites not EPs (blue) and without N-glycosylated sites (black). Unpaired Wilcoxon test was used to assess significance among groups with Bonferroni correction for multiple comparisons. **H)** Boxplot showing the highest APR strength (TANGO score) in β-sandwich domains for the same categories as G. Significance was assessed as in G. **I)** Schematic model of the quality control system of glycoproteins. **J)** Fraction of UGGT substrates that have an N-glycosite in an EP (yellow) or other N-glycosites (blue), as compared to all glycoproteins.

The high relative contact order observed in domains bearing protected APRs could be indicative of an enrichment for a specific fold topology, as most folds have lower contact orders than these particular domains (**Figure 5D**). To investigate this, we looked at the relative frequencies of protected APRs in different CATH architectures. As background, we used all SP protein domains with at least one APR. Interestingly, there was an underrepresentation of protected APRs in architectures of the class ‘Mainly alpha’, while architectures with more β-sheet content were more abundant (**Figure 5E**). In particular, the ‘CATH 2.60’ architecture (β-sandwich) was highly enriched and included the majority of N-glycosites at EP, which are distributed throughout the entire fold (**Figure 5E and 5F**). The β-sandwich architecture is characterised by two opposing antiparallel β-sheets and span a large number of fold superfamilies, including the immunoglobulin-like fold, and it has been linked to many neurodegenerative diseases associated with the formation of protein aggregates [51, 52]. Moreover, β-sandwich domains are frequently organised in linear arrays within multidomain proteins, which have a higher risk of forming domain-swapped misfolded species [53]. A deeper analysis of β-sandwich domains showed that those with N-glycosites at EPs have a stronger and higher number of APRs than the rest of β-sandwich domains, including other domains that are also N-glycosylated (**Figures 5G and 5H**). Furthermore, β-sandwich domains containing APRs protected by N-glycans are found in larger multidomain proteins, with, on average, 5 β-sandwich domains per protein (**Supplementary Figures 15E and 15F**).

N-glycosylation plays a crucial role in glycoprotein quality control (**Figure 5I**), as it acts as the attachment site for the ER soluble and membrane-bound lectin chaperones calreticulin and calnexin [30]. These chaperones have been shown to direct protein folding, reduce aggregation, retain misfolded or immature proteins within the ER and target aberrant proteins for degradation [54]. Upon release from the lectin chaperones, correctly folded proteins are transported to the Golgi apparatus. However, nascent chains that are not properly folded can be recognised by the protein folding sensor UDP-glucose:glycoprotein glucosyltransferase (UGGT) and then directed for rebinding to the lectin chaperones [55]. In other words, UGGT substrates are prone to misfold and require multiple rounds of chaperone binding. Recently, Adams *et al* [56] identified 71 *bona fide* human UGGT substrates using quantitative proteomics in HEK293 cells. Interestingly, proteins containing N-glycosites in EPs are significantly enriched in UGGT substrates when compared to other glycoproteins (**Figure 5J**).

Our findings show that the protection of APRs through N-glycans is linked to biophysical properties that challenge protein folding, such as structural complexity, a higher number of APRs and higher aggregation propensities. Moreover, this protection is enriched in UGGT substrates, which require multiple rounds of chaperone association to reach their native conformations. Therefore, it appears that these sites are strongly correlated with folding challenges, consistent with the idea that N-glycans mitigate aggregation prior to folding. In addition, the fact that the majority of domains that require this anti-aggregation mechanism have the same topology suggests that their folding landscapes, populated by similar folding intermediates [57], might have co-evolved together with N-glycosylation to avoid aggregation.

### Absence of N-glycosylation *in vivo* specifically increases protein aggregation

If N-glycosylation is indeed a general evolutionary measure against protein aggregation, its inhibition should affect protein solubility across proteomes. Indeed, in animal and plant cells, inhibition of N-glycosylation with tunicamycin leads to misfolding and aggregation inside the ER [58–60], triggering the unfolded protein response. To investigate which particular glycoproteins aggregate in the absence of N-glycosylation, we reanalysed a proteomics dataset from Sui *et al* [61]. In short, in this study they measured the changes in proteome solubility in the mouse Neuro2a cell line after treatment with six different stresses, including tunicamycin. Our analysis found that after treatment with tunicamycin, around 20% of the proteins identified with an N-glycosite at an EP are more insoluble (**Figure 6A and Supplementary Table 3**). Interestingly, in the majority of these aggregated proteins, the N-glycosite is located within a β-sandwich domain (**Supplementary Table 3**). In contrast, just 10% of proteins identified with N-glycosites that are not in EPs are more insoluble, suggesting that the absence of N-glycosylation alone has a smaller effect. However, due to the small number of proteins identified by the MS/MS, this difference was not statistically significant. Expectedly, an even smaller percentage of non-glycosylated proteins are found to be more insoluble after tunicamycin treatment. The same analysis was performed by looking at the solubility changes under the other five stresses. However, none of these affected the solubility of proteins identified with an N-glycosite in an EP (**Figure 6A**). The same was true when analysing proteins that are more soluble after each treatment (**Figure 6B**). Together, these results suggest that inhibiting N-glycosylation leads to a decrease in protein solubility, especially in proteins where N-glycosites are acting as aggregation gatekeepers (**Figure 6C**).

**Figure 6.**
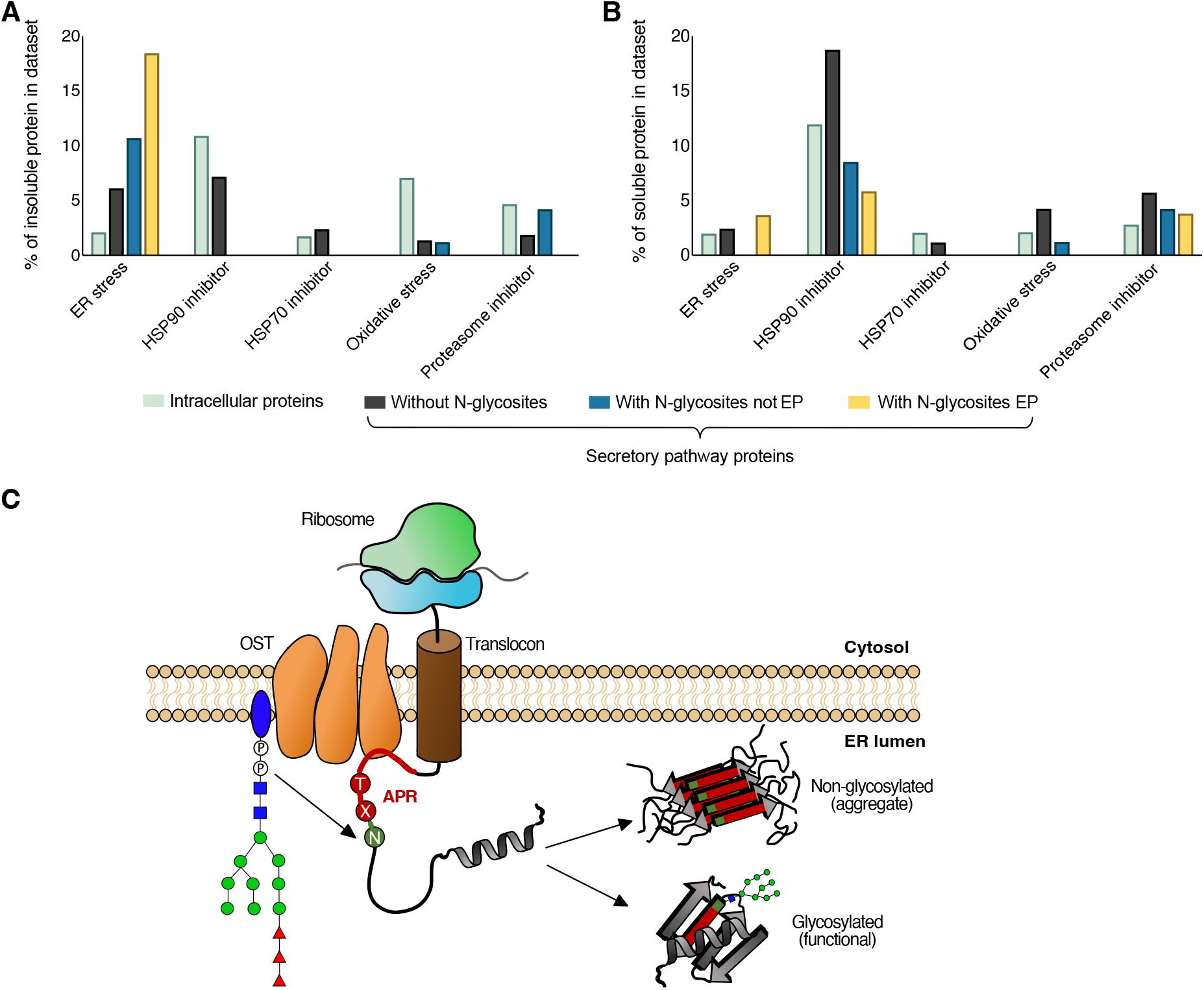
The absence of N-glycosylation *in vivo* specifically increases protein aggregation. **A)** Percentage of SP proteins that are enriched in the insoluble fraction in each group based on all proteins that are identified in the MS for that particular group. **B)** Percentage of SP proteins that are enriched in the soluble fraction in each group based on all proteins that are identified in the MS for that particular group. **C)** During translocation, the oligosaccharyltransferase (OST) can glycosylate proteins before these are folded. When N-glycans are attached at the flanks of an APR, they shield this region from aggregation, leading to a glycosylated functional protein. However, the absence of N-glycosylation, specifically at the flanks of an APR, can lead to the misfolding and aggregation of the affected proteins.

## DISCUSSION

Our work demonstrates that N-glycans are enriched, highly conserved and commonly replace unmodified gatekeeper residues in sequence segments with an intrinsic capacity to aggregate, here referred to as APRs, in nearly a thousand human proteins. In addition, we show that N-glycans suppress the aggregation of APRs *in vitro*, and that their inhibition in mouse Neuro2a cells leads to a specific aggregation of newly made proteins. Together, these findings suggest that, among its many molecular functions, N-glycosylation constitutes a functional mechanism directly dedicated to the control of protein aggregation in higher eukaryotes.

Many studies have shown that N-glycosylation prevents the aggregation of glycoproteins in cells through diverse indirect molecular mechanisms. For example, N-glycans can affect the folding process by restricting the conformational entropy of the unfolded protein and stabilising specific secondary structural elements, preventing the formation of folding intermediates prone to aggregate [54, 62]. Moreover, the association of glycoproteins with ER lectin chaperones increases folding efficiency while decreasing aggregation propensity [54]. Direct inhibition of aggregation by N-glycans has also been described, particularly in recombinant therapeutic proteins [63]. Indeed, for the production of therapeutic antibodies, such as bevacizumab, N-glycosylation sites have been engineered near APRs to mitigate aggregation [64]. However, the conditions in which biotherapeutics are produced are far from those found in cells, as often these proteins are manufactured and stored at very high concentrations for extended periods of time. Instead, our work points to a widely conserved cellular strategy in which N-glycans directly hinder the formation of aggregates during folding under physiological conditions.

A surprising result from our computational analysis is that only N-glycans located in the N-terminal flanks of APRs are under selection and share similar features to unmodified gatekeeper residues (**Figures 2 and 3**). However, placing N-glycans on either side of APRs *in vitro* strongly suppresses their aggregation (**Figure 4**). Since most N-glycans are co-translationally attached to proteins by STT3A, it appears possible that the preferential addition of this modification to the N-terminal flanks is coupled with translation. It has been proposed that the initiation of aggregation may occur within polysomes, where identical unfolded nascent chains reach high local concentrations [9, 65]. Under this framework, N-glycosylating the N-terminal side of an APR will immediately shield it from potential co-translational non-native interactions, including aggregation, as this side is translated before the rest of the APR sequence. Consistent with this hypothesis, the analysis of previously identified human STT3B-dependent sites [66] – specifically only attached post-translationally – showed a significant underrepresentation in EPs (**Supplementary Figure 16**). Moreover, overexpression of STT3B only partially rescues STT3A-deficient cells, despite STT3B acting downstream of STT3A, which enables it to glycosylate sites missed by STT3A [67]. Eukaryotic species lacking the STT3A ortholog, such as *Saccharomyces cerevisiae*, can only perform N-glycosylation post-translationally [68]. Indeed, unlike the other eukaryotic species analysed, the relative frequencies of glycosylated sites in EPs of yeast proteins were found to be underrepresented (**Figure 2F**). Despite all this circumstantial evidence, future studies are required to determine if N-glycosylation is specifically supressing aggregation during translation.

One question remains: why is N-glycosylation the only modification found to broadly act as an aggregation gatekeeper? Although we do not rule out that other PTM types not investigated here may act as gatekeepers, the answer probably again lies in the co-translational nature of this modification. Firstly, post-translational modifications require acceptor sites to be accessible to the modifying enzyme, precluding regions that are buried or structurally too rigid when the protein is folded, such as APRs and their GRs. Indeed, the placement of N-glycosylation in bacteria, which takes place post-translationally, is restricted only to flexible segments [69]. Therefore, coupling N-glycosylation with folding increases the number of sites that can be modified. Secondly, during folding, APRs are exposed and at risk of aggregation. Consequently, protein folding exerts a dual selection pressure on the glycosylation process [37]. On the one hand, sites that destabilise the native structure are under negative selection [70], while sites that optimise folding, in this case, by reducing aggregation, are under positive selection and are likely to become essential (**Figure 2E**). An additional consequence of this shift in the temporal sequence of maturation events has been the co-evolution of N-glycans with the ER chaperone machinery, leading to a very specific QC system for secretory and membrane glycoproteins [37]. Recently, a similar co-adaptation process was described between chaperone specificity and protein composition to explain the preference of Hsp70 for positively charged residues in bacteria [15].

We found a higher proportion of aggregated proteins with N-glycans acting as gatekeeper residues compared to other glycoproteins after treatment of mouse Neuro2a cells with tunicamycin (**Figure 6A**). Interestingly, tunicamycin treatment has been extensively used as a model to mimic type-I congenital disorders of glycosylation (CDG-I) [71, 72]. These are a rare group of metabolic diseases that affect specific sugar transferases and enzymes involved in the synthesis and transfer of N-glycans, thus leading to the improper N-glycosylation of proteins, which causes various symptoms potentially affecting multiple organs [73, 74]. It has been reported that several CDG-I can lead to ER stress and activate the unfolded protein response due to misfolded hypoglycosylated proteins unable to leave the ER [75]. Based on our findings, we hypothesize that the formation of protein aggregates resulting from a loss of N-glycans may provide an additional molecular cause of ER stress in CDGs and may contribute to the pathomechanism of these disorders. Future efforts should be made to determine if there is a direct relationship between these genetic disorders and protein aggregation.

## METHODS

### Human proteome dataset

The human proteome was obtained from UniProtKB/Swiss-Prot database (reference proteome UP000005640; release 2022_02). The dataset contains 19,379 proteins, after excluding sequences with nonstandard amino acids (e.g., selenocysteine), sequences with <25 amino acids and those with >10,000 residues and after filtering at 90% sequence identity using the CD-hit algorithm [76]. Signal peptides and transmembrane domains were identified using deepTMHMM [77] and removed from the analyses to avoid biases. In addition, deepTMHMM provides information on the overall topology of the protein. Experimentally annotated protein PTM sites were obtained from dbPTM [26] and from the GlcNAcAtlas [27], and were mapped to the proteome. Only those PTM types with more than 1,200 sites were retained.

Information on protein subcellular location was extracted from UniProt. Proteins known to reside in the endoplasmic reticulum, Golgi apparatus, cell membrane or extracellular space were labelled as part of the secretory pathway (SP). On the other hand, proteins known to reside in the cytoplasm, nucleus or mitochondria were labelled as part of the non-secretory pathway (non-SP). Proteins labelled both as SP and non-SP were excluded from further analyses.

Structural information was added to the dataset for each protein using the structures from the AlphaFold database [78, 79]. Absolute solvent accessibility values were calculated with DSSP based on these structures [80, 81]. Then, the relative solvent accessibility (RSA) values were calculated by dividing the absolute solvent accessibility values by residue-specific maximal accessibility values, as extracted from Tien *et al* [82]. Residues with RSA values < 0.2 were labelled as buried. Intrinsically disordered regions (IDRs) were identified using the pLDDT score provided in the AlphaFold models, as regions with low confidence scores have been shown to overlap largely with IDRs [83]. Residues with pLDDT scores < 50 were labelled as disordered.

### Protein aggregation prediction

Aggregation-prone regions (APRs) were predicted computationally using TANGO [1] at physiological conditions (pH at 7.5, temperature at 298 K, protein concentration at 1 mM, and ionic strength at 0.15 M). In this study, APRs are defined as segments between 5 and 15 amino acids in length, each with an aggregation score of at least 10. Gatekeeping regions (GRs) are defined as the three residues immediately downstream and upstream of APRs. All other residues are defined as distal regions (DRs).

APRs were also identified with CamSol [32]. CamSol calculates an intrinsic solubility profile where regions with a score higher than 1 are highly soluble, while scores smaller than −1 are poorly soluble (aggregation-prone). CamSol APRs are defined as segments between 5 and 15 amino acids in length, each with a solubility score smaller than −1. GRs and DRs are defined in the same way as above.

### Identification of sequons

All human proteins were scanned for N-glycosylation sequons (Asn-X-Thr/Ser, where X ≠ Pro). Sequons known to be glycosylated based on dbPTM annotations were labelled as “SP glycosylated”. Sequons in proteins from the secretory pathway without dbPTM annotations were labelled as “SP non-glycosylated”. Sequons in proteins that do not follow the secretory pathway, and thus cannot be glycosylated, were labelled as “non-SP”.

### Relative frequency calculation

For all PTM types and sequons, the frequency in each region was calculated by taking all verified PTMs in APRs, GRs and DRs versus all sites that could receive a PTM in each region:

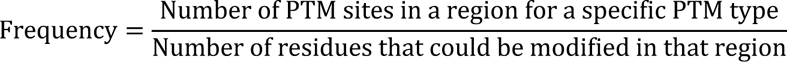

For example, for serine phosphorylation:

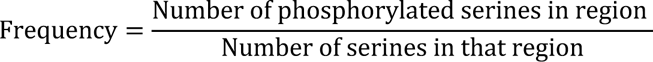

The relative frequency was obtained by dividing the frequency in each region by the overall frequency of that particular PTM (background). To avoid biases, only proteins that contain PTM sites are used as background.

### Eukaryotic proteome dataset

The proteomes of five other eukaryotic species were analysed in the same way as the human proteome and include representatives from the animal (*Mus musculus (UP000000589)*, *Drosophila melanogaster (UP000000803)* and *Caenorhabditis elegans (UP000001940)*), plant (*Arabidopsis thaliana (UP000006548)*) and fungal (*Saccharomyces cerevisiae (UP000002311)*) kingdom.

As for the human dataset, the frequencies of glycosylated and non-glycosylated sequons were determined for each eukaryotic species. However, since experimentally identified glycosylated sites for these organisms are scarce, all sequons in SP proteins were considered glycosylated (unless topological annotations by deepTMHMM [77] predicted the site to be facing the cytoplasm, where glycosylation does not occur).

### Sequon conservation analysis

The multiz100way [36] is a dataset containing multiple sequence alignments of 100 mammalian species to the human genome (hg38). Human N-glycosites were mapped to this dataset to calculate their conservation. A sequon is considered absent in a species (not conserved) if it deviates from the consensus sequence (N-X-T/S, where X ≠ P).

### Peptide set design

To construct a set of aggregating peptides with N-glycans at the flanks, APRs were selected from the human proteome containing an N-glycosylation site at the N-terminal or C-terminal flank. To facilitate accurate concentration determination of peptides through absorbance measurements at 280 nm, only APRs containing Trp and/or Tyr were considered. 20 APRs were synthesised and screened for ThT-binding kinetics, from which a final set of ten APRs was selected based on their kinetic profile. Five of these sequences had the N-glycosylation site in the N-terminal, while the other five were in the C-terminal. Seven variants for each peptide sequence were produced: non-modified (WT), GlcNAc, Man9 (Man_9_GlcNAc_2_), and each of the charged residues (D, E, K, and R). For two specific peptide sets, four more variants were produced: GlcNAc_2_, ManGlcNAc_2_ (Man), Man_3_GlcNAc_2_ (Man3), and Man_6_GlcNAc_2_ (Man6).

### Peptide aggregation kinetics

All peptides, except the GlcNAc_2_, Man, Man3 and Man9 variants, were synthesised in-house using an Intavis Multipep RSi solid-phase peptide synthesis robot. The complex glycoform peptide variants were ordered from Chemitope Glycopeptide. Stocks were then diluted to the appropriate peptide concentration in PBS with a final concentration of 5% DMSO. The concentration for each peptide set was selected based on their ThT-binding kinetic profile to have a lag phase shorter than 72 hours. TCEP (1 mM) was included in solutions of peptides containing cysteine or methionine residues to disrupt disulphide bond formation. For ThT- and pFTAA-binding kinetics, 10 μM ThT or 1 μM pFTAA were added to the peptide samples. Dye binding was measured over time through excitation at 440 nm and emission at 480 and 520 nm, for ThT and pFTAA, respectively, in a Fluostar OMEGA.

For Endo H treatment, endoglycosidase H (500 units, 1 μl, New England Biolabs, catalog no. P0702) was added to each of the SLNYLLYVSN peptide samples. Aggregation kinetics were measured over time as above.

### Endpoint solubility

For endpoint solubility concentrations, peptide preparations were left at room temperature for a week at an initial concentration equal to the one used in for aggregation kinetics. Peptides were subsequently subjected to ultracentrifugation at 100,000 g for 1h at 4°C. Supernatant concentrations were measured using RP-HPLC. Concentrations were measured with RP-HPLC instead of using absorbance measurements at 280 nm since it is more accurate for low concentrations.

### TEM imaging

Peptide solutions were incubated for a week at room temperature at the same concentrations of previous experiments. Suspensions (5 μL) of each peptide solution were added on 400-mesh carbon-coated copper grids, which were negatively stained using uranyl acetate. Grids were examined with a JEM-1400 120 kV transmission electron microscope.

### Machine learning

To predict which APRs are protected by N-glycans, a random forest classifier (randomForest R package, number of trees = 500, mtry = 3) was trained using several features of the APRs (relative solvent accessibility, length, number of unmodified gatekeepers, relative position within the domain, aggregation propensity, sequence disorder and number of cysteines) and of the protein domains bearing such APRs (contact order, length, number of APRs per 100 amino acids and number of disulphide bonds per 100 amino acids). Domain boundaries were extracted using CATH-Gene3D [48, 49]. The contact order for each domain was calculated as defined by Plaxco *et al.* [50]. The number of disulphide bonds was extracted from UniProt. Random undersampling and random oversampling (ROSE R package) were used to avoid biases due to class imbalance. Feature importance was evaluated with the Mean Decrease Accuracy plot, which indicates how much accuracy the model loses when excluding each variable.

### Statistics

GraphPad prism or R software were used to perform the different statistical tests. The tests used in each analysis are specified in the corresponding figure. *P*-values are represented as: * *P*-value ≤ 0.05, ** *P*-value ≤ 0.01, *** *P*-value ≤ 0.001.

### Visualisations

Visualisations were performed with GraphPad prism or custom R scripts using the packages ggplot2 [84] and ComplexHeatmap [85]. ChimeraX was used to visualize protein structures [86].

## ABBREVIATIONS

APR: Aggregation-prone region
PTM: Post-translational modification
GR: Gatekeeping region
DR: Distal region
SP: Secretory pathway
EP: Enriched position
OST: Oligosaccharyltransferase
ER: Endoplasmic reticulum
CDG: Congenital disorders of glycosylation

## ACKNOWLEDGEMENTS

The Switch Laboratory was supported by the Flanders Institute for Biotechnology (VIB, grant no. C0401 to FR and JS); the Fund for Scientific Research Flanders (FWO, project grant G053420N to JS; and Postdoctoral Fellowships 12P0919N and 12P0922N to NL, 12S3722N to BH and 1289023N to MPW); the Jaeken-Theunissen CDG Fund (to GM and MPW) and a CELSA fund grant (CELSA/21/027 to GM and MPW). The authors gratefully acknowledge the Electron Microscopy Platform of the VIB – KU Leuven Center for Brain & Disease Research for their support & assistance in this work.

## AUTHOR CONTRIBUTIONS

**FR, JS** and **NL** conceived and supervised this study. **RDR, FR, JS, NL, BH, MPW and GM** and designed experiments and *in silico* analyses. **RDR** performed *in vitro* experimental work, as well as all *in silico* analyses. **MDV** performed peptide synthesis. **RDR, FR and JS** and wrote the manuscript. All authors proofread and corrected the manuscript.

## REFERENCES

[1] Fernandez-Escamilla AM, Rousseau F, Schymkowitz J, Serrano L. Prediction of sequence-dependent and mutational effects on the aggregation of peptides and proteins. Nat Biotechnol. 2004;22:1302–6.

[2] Rousseau F, Serrano L, Schymkowitz JW. How evolutionary pressure against protein aggregation shaped chaperone specificity. J Mol Biol. 2006;355:1037–47.

[3] Prabakaran R, Goel D, Kumar S, Gromiha MM. Aggregation prone regions in human proteome: Insights from large-scale data analyses. Proteins: Structure, Function, and Bioinformatics. 2017;85:1099–118.

[4] Tyedmers J, Mogk A, Bukau B. Cellular strategies for controlling protein aggregation. Nat Rev Mol Cell Biol. 2010;11:777–88.

[5] Saibil H. Chaperone machines for protein folding, unfolding and disaggregation. Nature reviews Molecular cell biology. 2013;14:630–42.

[6] Chiti F, Dobson CM. Protein Misfolding, Amyloid Formation, and Human Disease: A Summary of Progress Over the Last Decade. Annu Rev Biochem. 2017;86:27–68.

[7] Iadanza MG, Jackson MP, Hewitt EW, Ranson NA, Radford SE. A new era for understanding amyloid structures and disease. Nat Rev Mol Cell Biol. 2018;19:755–73.

[8] Langenberg T, Gallardo R, van der Kant R, Louros N, Michiels E, Duran-Romana R, et al. Thermodynamic and Evolutionary Coupling between the Native and Amyloid State of Globular Proteins. Cell reports. 2020;31:107512.

[9] Houben B, Rousseau F, Schymkowitz J. Protein structure and aggregation: a marriage of necessity ruled by aggregation gatekeepers. Trends Biochem Sci. 2022;47:194–205.

[10] Monsellier E, Ramazzotti M, Taddei N, Chiti F. Aggregation Propensity of the Human Proteome. Plos Computational Biology. 2008;4.

[11] Buell AK, Tartaglia GG, Birkett NR, Waudby CA, Vendruscolo M, Salvatella X, et al. Position-Dependent Electrostatic Protection against Protein Aggregation. Chembiochem. 2009;10:1309–12.

[12] Markiewicz BN, Oyola R, Du D, Gai F. Aggregation Gatekeeper and Controlled Assembly of Trpzip β-Hairpins. Biochemistry. 2014;53:1146–54.

[13] Sant’Anna R, Braga C, Varejão N, Pimenta KM, Graña-Montes R, Alves A, et al. The Importance of a Gatekeeper Residue on the Aggregation of Transthyretin: IMPLICATIONS FOR TRANSTHYRETIN-RELATED AMYLOIDOSES. Journal of Biological Chemistry. 2014;289:28324–37.

[14] Beerten J, Jonckheere W, Rudyak S, Xu J, Wilkinson H, De Smet F, et al. Aggregation gatekeepers modulate protein homeostasis of aggregating sequences and affect bacterial fitness. Protein engineering, design & selection : PEDS. 2012;25:357–66.

[15] Houben B, Michiels E, Ramakers M, Konstantoulea K, Louros N, Verniers J, et al. Autonomous aggregation suppression by acidic residues explains why chaperones favour basic residues. EMBO J. 2020;39:e102864.

[16] De Baets G, Van Durme J, Rousseau F, Schymkowitz J. A genome-wide sequence-structure analysis suggests aggregation gatekeepers constitute an evolutionary constrained functional class. J Mol Biol. 2014;426:2405–12.

[17] De Baets G, Van Doorn L, Rousseau F, Schymkowitz J. Increased Aggregation Is More Frequently Associated to Human Disease-Associated Mutations Than to Neutral Polymorphisms. PLoS Comput Biol. 2015;11:e1004374.

[18] Schaffert LN, Carter WG. Do Post-Translational Modifications Influence Protein Aggregation in Neurodegenerative Diseases: A Systematic Review. Brain Sci. 2020;10.

[19] Alquezar C, Arya S, Kao AW. Tau post-translational modifications: dynamic transformers of tau function, degradation, and aggregation. Frontiers in neurology. 2021;11:595532.

[20] Barrett PJ, Timothy Greenamyre J. Post-translational modification of alpha-synuclein in Parkinson’s disease. Brain research. 2015;1628:247–53.

[21] Rezaei-Ghaleh N, Kumar S, Walter J, Zweckstetter M. Phosphorylation interferes with maturation of amyloid-β fibrillar structure in the N terminus. Journal of Biological Chemistry. 2016;291:16059–67.

[22] Gong C-X, Liu F, Grundke-Iqbal I, Iqbal K. Post-translational modifications of tau protein in Alzheimer’s disease. Journal of neural transmission. 2005;112:813–38.

[23] Ryan P, Xu M, Davey AK, Danon JJ, Mellick GD, Kassiou M, et al. O-GlcNAc modification protects against protein misfolding and aggregation in neurodegenerative disease. ACS chemical neuroscience. 2019;10:2209–21.

[24] Martinez MR, Dias TB, Natov PS, Zachara NE. Stress-induced O-GlcNAcylation: an adaptive process of injured cells. Biochemical Society Transactions. 2017;45:237–49.

[25] Pearlman SM, Serber Z, Ferrell JE. A mechanism for the evolution of phosphorylation sites. Cell. 2011;147:934–46.

[26] Li Z, Li S, Luo M, Jhong J-H, Li W, Yao L, et al. dbPTM in 2022: an updated database for exploring regulatory networks and functional associations of protein post-translational modifications. Nucleic acids research. 2022;50:D471–D9.

[27] Ma J, Li Y, Hou C, Wu C. O-GlcNAcAtlas: A database of experimentally identified O-GlcNAc sites and proteins. Glycobiology. 2021;31:719–23.

[28] Pang CNI, Hayen A, Wilkins MR. Surface Accessibility of Protein Post-Translational Modifications. Journal of Proteome Research. 2007;6:1833–45.

[29] Bludau I, Willems S, Zeng W-F, Strauss MT, Hansen FM, Tanzer MC, et al. The structural context of posttranslational modifications at a proteome-wide scale. PLOS Biology. 2022;20:e3001636.

[30] Helenius A, Aebi M. Roles of N-linked glycans in the endoplasmic reticulum. Annual review of biochemistry. 2004;73:1019–49.

[31] Varki A. Biological roles of glycans. Glycobiology. 2017;27:3–49.

[32] Sormanni P, Aprile FA, Vendruscolo M. The CamSol method of rational design of protein mutants with enhanced solubility. Journal of molecular biology. 2015;427:478–90.

[33] Malaby HL, Kobertz WR. The middle X residue influences cotranslational N-glycosylation consensus site skipping. Biochemistry. 2014;53:4884–93.

[34] Igura M, Kohda D. Quantitative assessment of the preferences for the amino acid residues flanking archaeal N-linked glycosylation sites. Glycobiology. 2011;21:575–83.

[35] Huang Y-W, Yang H-I, Wu Y-T, Hsu T-L, Lin T-W, Kelly JW, et al. Residues comprising the enhanced aromatic sequon influence protein N-glycosylation efficiency. Journal of the American Chemical Society. 2017;139:12947–55.

[36] Blanchette M, Kent WJ, Riemer C, Elnitski L, Smit AF, Roskin KM, et al. Aligning multiple genomic sequences with the threaded blockset aligner. Genome Res. 2004;14:708–15.

[37] Aebi M. N-linked protein glycosylation in the ER. Biochimica et Biophysica Acta (BBA)-Molecular Cell Research. 2013;1833:2430–7.

[38] Lombard J. The multiple evolutionary origins of the eukaryotic N-glycosylation pathway. Biology direct. 2016;11:1–31.

[39] Reumers J, Maurer-Stroh S, Schymkowitz J, Rousseau Fd. Protein sequences encode safeguards against aggregation. Human Mutation. 2009;30:431–7.

[40] Law RH, Zhang Q, McGowan S, Buckle AM, Silverman GA, Wong W, et al. An overview of the serpin superfamily. Genome Biol. 2006;7:1–11.

[41] Spence MA, Mortimer MD, Buckle AM, Minh BQ, Jackson CJ. A comprehensive phylogenetic analysis of the serpin superfamily. Molecular Biology and Evolution. 2021;38:2915–29.

[42] Stanley P. Golgi glycosylation. Cold Spring Harbor perspectives in biology. 2011;3:a005199.

[43] Cherepanova NA, Venev SV, Leszyk JD, Shaffer SA, Gilmore R. Quantitative glycoproteomics reveals new classes of STT3A-and STT3B-dependent N-glycosylation sites. Journal of Cell Biology. 2019;218:2782–96.

[44] Nakamura H, Kiyoshi M, Anraku M, Hashii N, Oda-Ueda N, Ueda T, et al. Glycosylation decreases aggregation and immunogenicity of adalimumab Fab secreted from Pichia pastoris. The Journal of Biochemistry. 2021;169:435–43.

[45] Solá RJ, Griebenow K. Effects of glycosylation on the stability of protein pharmaceuticals. Journal of pharmaceutical sciences. 2009;98:1223–45.

[46] Mason PE, Lerbret A, Saboungi ML, Neilson GW, Dempsey CE, Brady JW. Glucose interactions with a model peptide. Proteins: Structure, Function, and Bioinformatics. 2011;79:2224–32.

[47] Han J-H, Batey S, Nickson AA, Teichmann SA, Clarke J. The folding and evolution of multidomain proteins. Nature Reviews Molecular Cell Biology. 2007;8:319–30.

[48] Sillitoe I, Bordin N, Dawson N, Waman VP, Ashford P, Scholes HM, et al. CATH: increased structural coverage of functional space. Nucleic acids research. 2021;49:D266–D73.

[49] Lewis TE, Sillitoe I, Dawson N, Lam SD, Clarke T, Lee D, et al. Gene3D: extensive prediction of globular domains in proteins. Nucleic acids research. 2018;46:D435–D9.

[50] Plaxco KW, Simons KT, Baker D. Contact order, transition state placement and the refolding rates of single domain proteins. J Mol Biol. 1998;277:985–94.

[51] Merlini G, Comenzo RL, Seldin DC, Wechalekar A, Gertz MA. Immunoglobulin light chain amyloidosis. Expert review of hematology. 2014;7:143–56.

[52] Grad LI, Fernando SM, Cashman NR. From molecule to molecule and cell to cell: prion-like mechanisms in amyotrophic lateral sclerosis. Neurobiology of disease. 2015;77:257–65.

[53] Borgia A, Kemplen KR, Borgia MB, Soranno A, Shammas S, Wunderlich B, et al. Transient misfolding dominates multidomain protein folding. Nature communications. 2015;6:8861.

[54] Hebert DN, Lamriben L, Powers ET, Kelly JW. The intrinsic and extrinsic effects of N-linked glycans on glycoproteostasis. Nature chemical biology. 2014;10:902–10.

[55] Sousa M, Parodi AJ. The molecular basis for the recognition of misfolded glycoproteins by the UDP-Glc: glycoprotein glucosyltransferase. The EMBO journal. 1995;14:4196–203.

[56] Adams BM, Canniff NP, Guay KP, Larsen ISB, Hebert DN. Quantitative glycoproteomics reveals cellular substrate selectivity of the ER protein quality control sensors UGGT1 and UGGT2. Elife. 2020;9:e63997.

[57] Neudecker P, Robustelli P, Cavalli A, Walsh P, Lundstrom P, Zarrine-Afsar A, et al. Structure of an intermediate state in protein folding and aggregation. Science. 2012;336:362–6.

[58] Tkacz JS, Lampen JO. Tunicamycin inhibition of polyisoprenyl N-acetylglucosaminyl pyrophosphate formation in calf-liver microsomes. Biochem Biophys Res Commun. 1975;65:248–57.

[59] Marquardt T, Helenius A. Misfolding and aggregation of newly synthesized proteins in the endoplasmic reticulum. The Journal of cell biology. 1992;117:505–13.

[60] Sparvoli F, Faoro F, Daminati MG, Ceriotti A, Bollini R. Misfolding and aggregation of vacuolar glycoproteins in plant cells. The Plant Journal. 2000;24:825–36.

[61] Sui X, Pires DEV, Ormsby AR, Cox D, Nie S, Vecchi G, et al. Widespread remodeling of proteome solubility in response to different protein homeostasis stresses. Proc Natl Acad Sci U S A. 2020;117:2422–31.

[62] Caramelo JJ, Parodi AJ. A sweet code for glycoprotein folding. FEBS letters. 2015;589:3379–87.

[63] Zhou Q, Qiu H. The mechanistic impact of N-glycosylation on stability, pharmacokinetics, and immunogenicity of therapeutic proteins. Journal of pharmaceutical sciences. 2019;108:1366–77.

[64] Courtois F, Agrawal NJ, Lauer TM, Trout BL. Rational design of therapeutic mAbs against aggregation through protein engineering and incorporation of glycosylation motifs applied to bevacizumab. MAbs. 2016;8:99–112.

[65] Brandt F, Etchells SA, Ortiz JO, Elcock AH, Hartl FU, Baumeister W. The native 3D organization of bacterial polysomes. Cell. 2009;136:261–71.

[66] Shrimal S, Trueman SF, Gilmore R. Extreme C-terminal sites are posttranslocationally glycosylated by the STT3B isoform of the OST. Journal of Cell Biology. 2013;201:81–95.

[67] Shrimal S, Ng BG, Losfeld M-E, Gilmore R, Freeze HH. Mutations in STT3A and STT3B cause two congenital disorders of glycosylation. Human molecular genetics. 2013;22:4638–45.

[68] Shrimal S, Cherepanova NA, Mandon EC, Venev SV, Gilmore R. Asparagine-linked glycosylation is not directly coupled to protein translocation across the endoplasmic reticulum in Saccharomyces cerevisiae. Molecular biology of the cell. 2019;30:2626–38.

[69] Lizak C, Gerber S, Numao S, Aebi M, Locher KP. X-ray structure of a bacterial oligosaccharyltransferase. Nature. 2011;474:350–5.

[70] Medus ML, Gomez GE, Zacchi LF, Couto PM, Labriola CA, Labanda MS, et al. N-glycosylation triggers a dual selection pressure in eukaryotic secretory proteins. Sci Rep-Uk. 2017;7:8788.

[71] Rita Lecca M, Wagner U, Patrignani A, Berger EG, Hennet T. Genome-wide analysis of the unfolded protein response in fibroblasts from congenital disorders of glycosylation type-I patients. The FASEB journal. 2005;19:1–21.

[72] de Haas P, de Jonge MI, Koenen HJ, Joosten B, Janssen MC, de Boer L, et al. Evaluation of Cell Models to Study Monocyte Functions in PMM2 Congenital Disorders of Glycosylation. Frontiers in Immunology. 2022;13.

[73] Wilson MP, Matthijs G. The evolving genetic landscape of congenital disorders of glycosylation. Biochimica et Biophysica Acta (BBA)-General Subjects. 2021;1865:129976.

[74] Yuste-Checa P, Vega AI, Martín-Higueras C, Medrano C, Gámez A, Desviat LR, et al. DPAGT1-CDG: Functional analysis of disease-causing pathogenic mutations and role of endoplasmic reticulum stress. PLoS One. 2017;12:e0179456.

[75] Sun L, Zhao Y, Zhou K, Freeze HH, Zhang Y-w, Xu H. Insufficient ER-stress response causes selective mouse cerebellar granule cell degeneration resembling that seen in congenital disorders of glycosylation. Molecular brain. 2013;6:1–8.

[76] Fu L, Niu B, Zhu Z, Wu S, Li W. CD-HIT: accelerated for clustering the next-generation sequencing data. Bioinformatics. 2012;28:3150–2.

[77] Hallgren J, Tsirigos KD, Pedersen MD, Armenteros JJA, Marcatili P, Nielsen H, et al. DeepTMHMM predicts alpha and beta transmembrane proteins using deep neural networks. bioRxiv. 2022.

[78] Jumper J, Evans R, Pritzel A, Green T, Figurnov M, Ronneberger O, et al. Highly accurate protein structure prediction with AlphaFold. Nature. 2021;596:583–9.

[79] Varadi M, Anyango S, Deshpande M, Nair S, Natassia C, Yordanova G, et al. AlphaFold Protein Structure Database: massively expanding the structural coverage of protein-sequence space with high-accuracy models. Nucleic Acids Res. 2022;50:D439–D44.

[80] Kabsch W, Sander C. Dictionary of protein secondary structure: pattern recognition of hydrogen-bonded and geometrical features. Biopolymers. 1983;22:2577–637.

[81] Joosten RP, te Beek TA, Krieger E, Hekkelman ML, Hooft RW, Schneider R, et al. A series of PDB related databases for everyday needs. Nucleic Acids Res. 2011;39:D411–9.

[82] Tien MZ, Meyer AG, Sydykova DK, Spielman SJ, Wilke CO. Maximum allowed solvent accessibilites of residues in proteins. PloS one. 2013;8:e80635.

[83] Ruff KM, Pappu RV. AlphaFold and implications for intrinsically disordered proteins. Journal of Molecular Biology. 2021;433:167208.

[84] Wickham H. Data analysis. ggplot2: Springer; 2016. p. 189–201.

[85] Gu Z, Eils R, Schlesner M. Complex heatmaps reveal patterns and correlations in multidimensional genomic data. Bioinformatics. 2016;32:2847–9.

[86] Goddard TD, Huang CC, Meng EC, Pettersen EF, Couch GS, Morris JH, et al. UCSF ChimeraX: Meeting modern challenges in visualization and analysis. Protein Science. 2018;27:14–25.

