## Supplementary materials for "N-glycosylation as a eukaryotic protective mechanism against protein aggregation"

#### Supplementary table 1

List of all PTM-types used for the enrichment analysis.

| PTM-type | Total occurrences | Residue | Source |
|---|---|---|---|
| Phosphoserine | 244,913 | Ser | dbPTM |
| Phosphothreonine | 109,301 | Thr | dbPTM |
| Lysine ubiquitination | 59,712 | Lys | dbPTM |
| Phosphotyrosine | 48,576 | Tyr | dbPTM |
| Lysine acetylation | 38,440 | Lys | dbPTM |
| N-Glycosylation | 15,047 | Asn | dbPTM |
| Serine O-GlcNAcylation | 7,122 | Ser | GlcNAcAtlas |
| Threonine O-Glycosylation | 6,341 | Thr | dbPTM |
| Serine O-Glycosylation | 6,061 | Ser | dbPTM |
| Arginine methylation | 5,614 | Arg | dbPTM |
| SUMOylation | 5,430 | Lys | dbPTM |
| Malonylation | 5,282 | Lys | dbPTM |
| Sulfoxidation | 4,973 | Met | dbPTM |
| Threonine O-GlcNAcylation | 4,578 | Thr | GlcNAcAtlas |
| Lysine methylation | 2,412 | Lys | dbPTM |
| Glutathionylation | 1,668 | Cys | dbPTM |
| Succinylation | 1,495 | Lys | dbPTM |

#### Supplementary table 2

List of all aggregation peptide cores used in this study.

| Peptide sequence | UniProt ID | Gene | APR position | TANGO score | Concentration (µM) | Side |
|---|---|---|---|---|---|---|
| SLNYLLYVSN | Q04912 | MST1R | 479 | 57.1 | 50 | C-ter |
| NASYILIR | Q76LX8 | ADAMTS13 | 1354 | 42.9 | 200 | N-ter |
| NLTTLTFWG | Q9NZ08 | ERAP1 | 70 | 33.4 | 50 | N-ter |
| NISCLWVFK | P36888 | FLT3 | 100 | 66.8 | 50 | N-ter |
| DFYICVN | Q14627 | IL13RA2 | 209 | 66.0 | 50 | C-ter |
| NFTYIID | P48551 | IFNAR2 | 192 | 83.5 | 250 | N-ter |
| NVSVYTVK | P02751 | FN1 | 1244 | 43.4 | 300 | N-ter |
| RYAVYWN | P52803 | EFNA5 | 31 | 78.4 | 300 | C-ter |
| QYVLWASN | Q53GD3 | SLC44A4 | 386 | 52.8 | 150 | C-ter |
| HSIYMFFN | P01137 | TGFB1 | 129 | 73.2 | 100 | C-ter |

#### Supplementary table 3

List of proteins that are more insoluble after tunicamycin treatment with N-glycosites in EPs.

| UniProt ID | Protein name | N-glycosite position | Side | In beta sandwich? |
|---|---|---|---|---|
| Q9R0E1 | Lysysl hydroxylase 3 | 66 | APR | No |
| P09055 | Integrin beta-1 | 366 | GR1 N-ter | Yes |
| Q61735 | Leukocyte surface antigen CD47 | 61 | GR2 N-ter | Yes |
| Q99KV1 | ER-associated DNAJ | 261 | GR1 N-ter | Yes |
| Q61739 | Integrin alpha-6 | 746 | GR2 N-ter | Yes |

### Supplementary figure 1

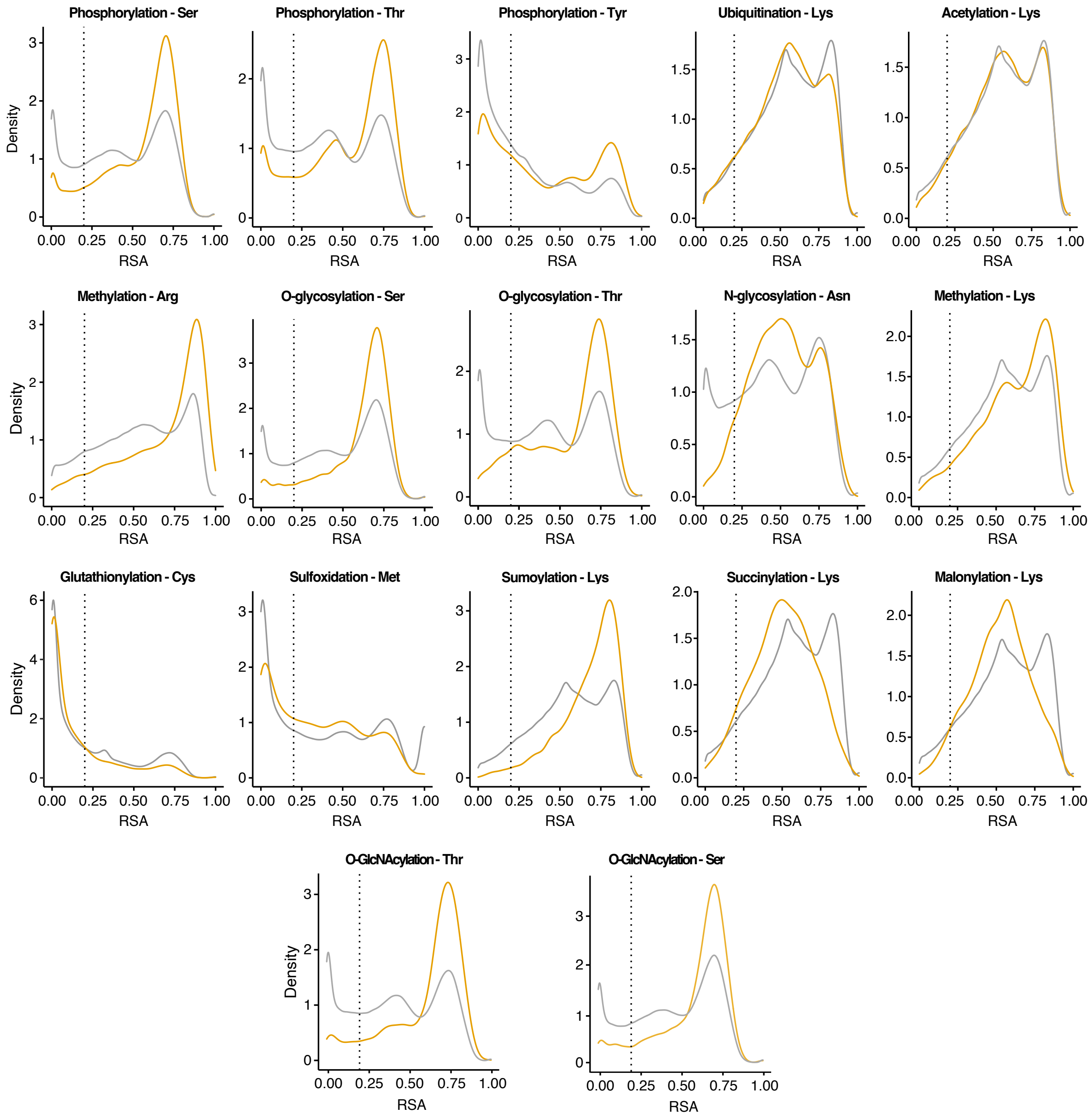

**Supplementary figure 1.** Distribution of relative solvent accessibility (RSA) values of experimentally determined sites (in gold) and amino acid residues (in grey) for all PTM-types analysed. RSA values <= 0.2 are considered completely buried (dot line).

#### Supplementary figure 2

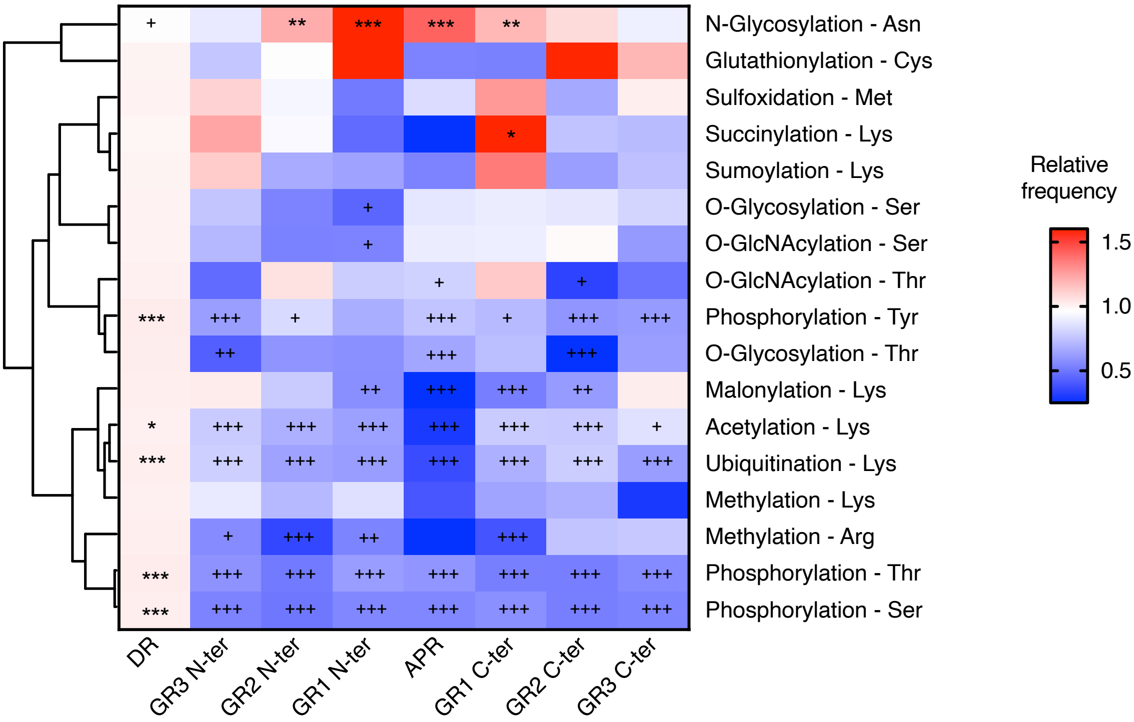

**Supplementary figure 2.** Heatmap showing the relative frequencies for each of the 15 types of PTMs on residues with an RSA > 0.2. Crosses (asterisks) indicate that a region has a significantly lower (higher) frequency compared to the background by Fisher exact test with FDR correction. Crosses (asterisks) at the top of the bar indicate that a region has a significantly lower (higher) frequency compared to the background by Fisher exact test with FDR correction. Rows are clustered based on Pearson correlation as distance measure.

#### Supplementary figure 3

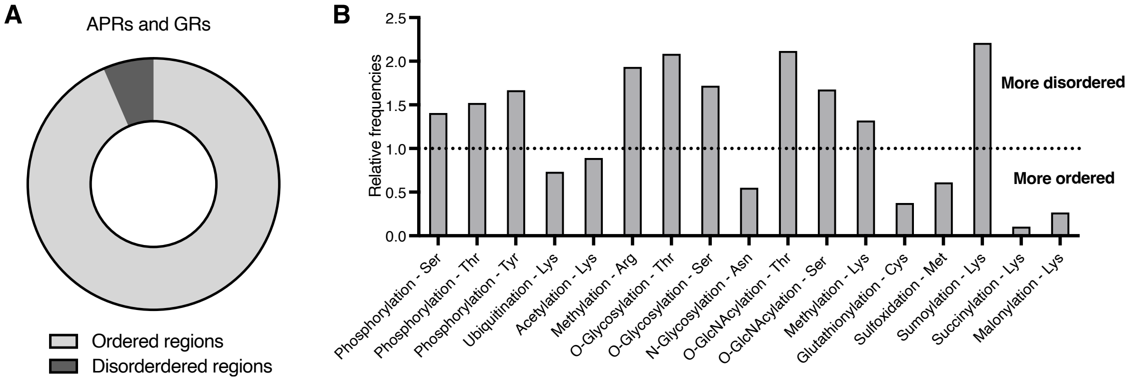

**Supplementary figure 3. A)** Fraction of predicted ordered (pLDDT >= 50) and disordered (pLDDT < 50) residues in APRs and GRs. **B)** Relative frequencies of different PTM-types in disordered regions. Values > 1 indicate that a PTM-type is found more often in disordered regions while values < 1 indicate that a PTM-type is found more often in ordered regions.

#### Supplementary figure 4

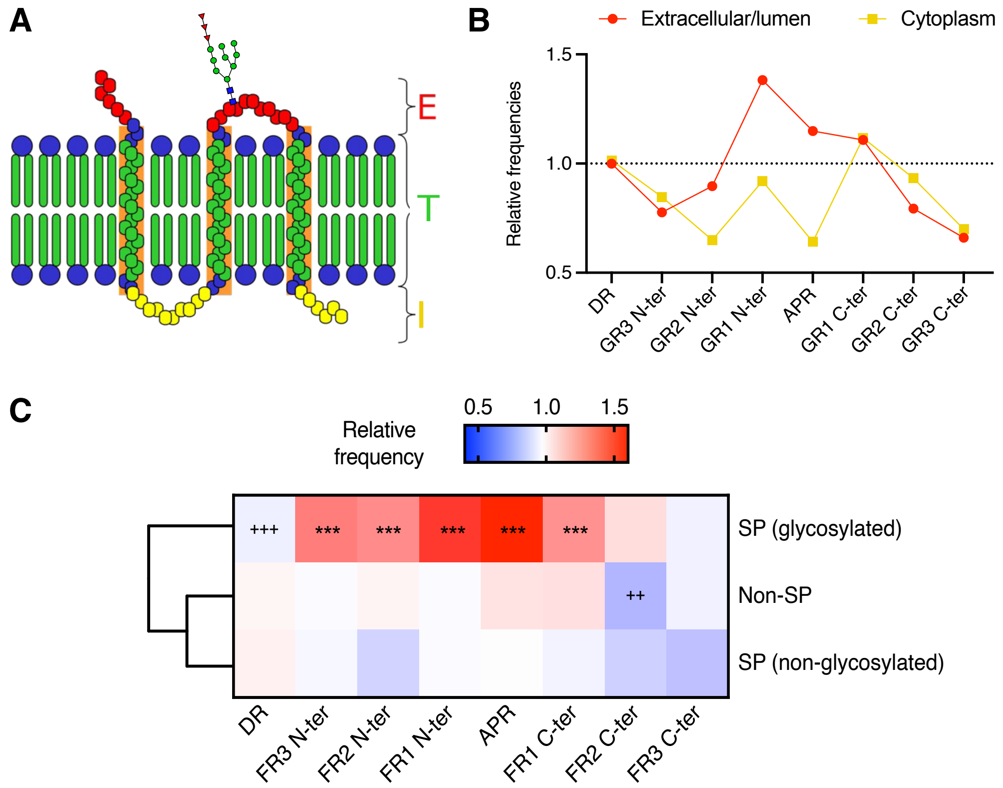

**Supplementary figure 4. A)** Transmembrane proteins have domains facing the extracellular/lumenal or the intracellular side of the cell, which can be predicted using deepTMHMM. Only sequons in domains facing the extracellular/lumenal side can get glycosylated. **B)** Relative frequencies of sequons in domains of transmembrane proteins facing the extracellular/lumenal (red) or intracellular side (yellow). **C)** Heatmap showing the relative frequencies of having a sequon in each region (columns) for different subsets of proteins (rows) for CamSol APRs. Crosses (asterisks) at the top of the bar indicate that a region has a significantly lower (higher) frequency compared to the background by Fisher exact test with FDR correction. Rows are clustered based on Pearson correlation as distance measure.

#### Supplementary figure 5

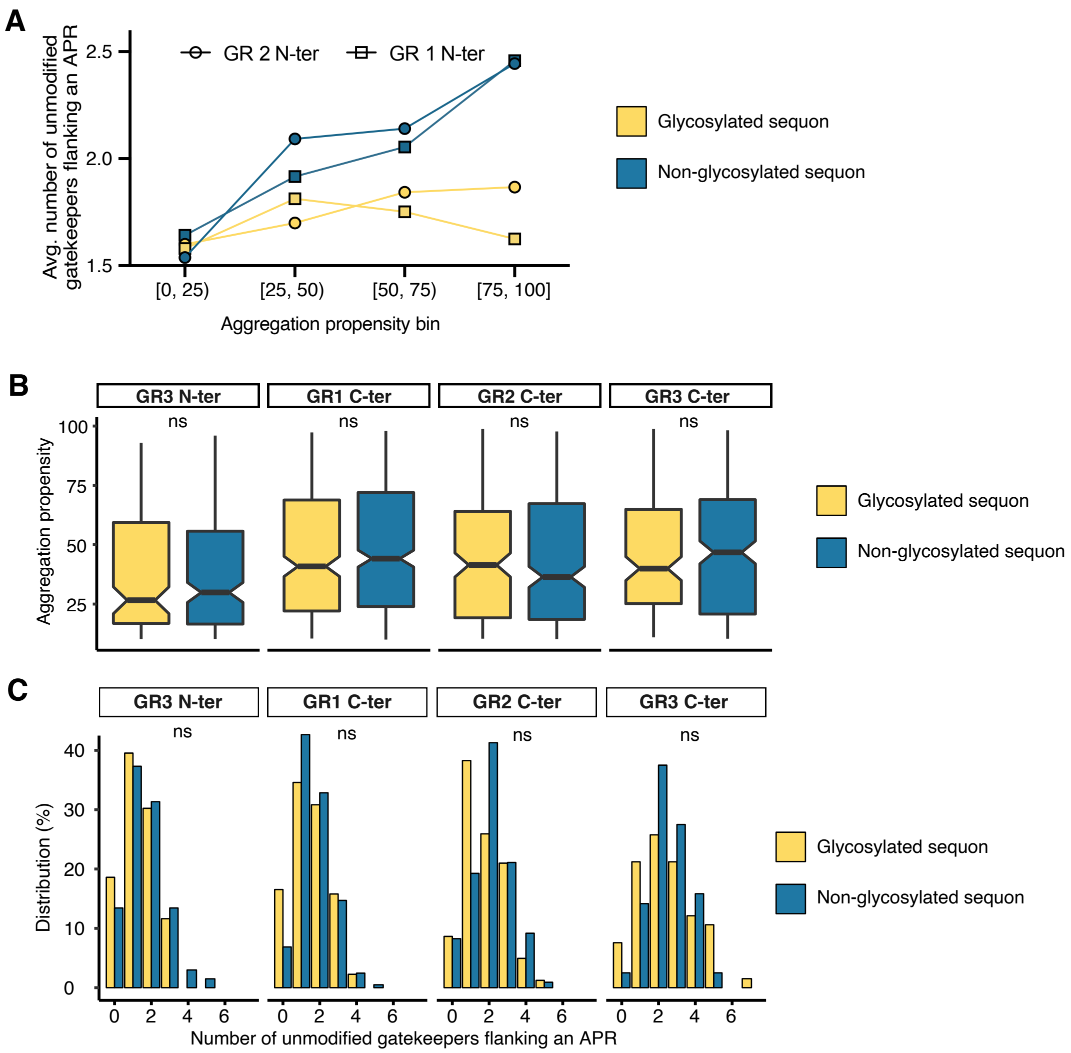

**Supplementary figure 5. A)** Average number of unmodified gatekeepers flanking APRs for glycosylated and non-glycosylated sequons in GR2 N-ter and GR1 N-ter. APRs are classified based on aggregation propensity (TANGO score) in four bins. **B)** Aggregation propensity (TANGO score) of APRs that have glycosylated sequons or have non-glycosylated sequons in GR3 N-ter, GR1 C-ter, GR2 C-ter and GR3 C-ter (non-EPs). Unpaired Wilcoxon test was used to assess significance between the two groups (glycosylated vs non-glycosylated sequons). **C)** Number of unmodified gatekeeping residues three positions upstream and three positions downstream for APRs with an aggregation propensity score >= 50. Unpaired Wilcoxon test was used to assess significance between the two groups.

#### Supplementary figure 6

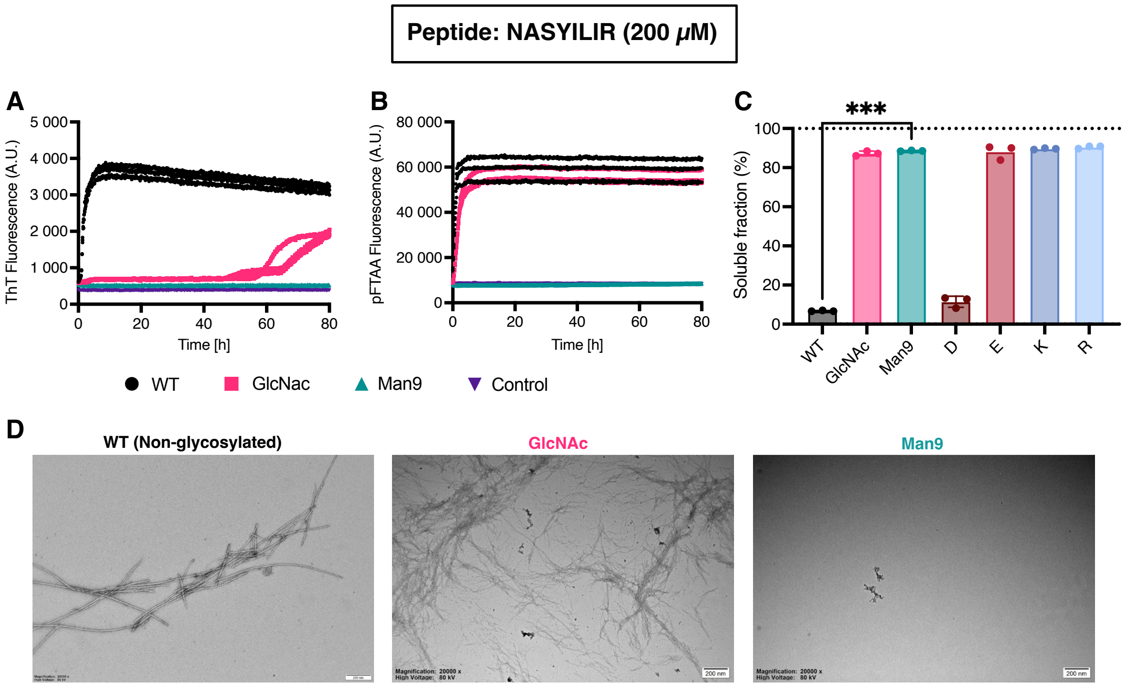

**Supplementary figure 6. A, B)** ThT binding (A) and pFTAA binding (B) kinetics. Fluorescence over time is shown for three independent repeats. Vehicle control fluorescence is shown in purple. **C)** Percentage of the concentration of peptide in the soluble fraction after ultracentrifugation (n=3). Unpaired t-test was used to assess significance of wt against Man9. **D)** TEM images after 7 days of incubation.

#### Supplementary figure 7

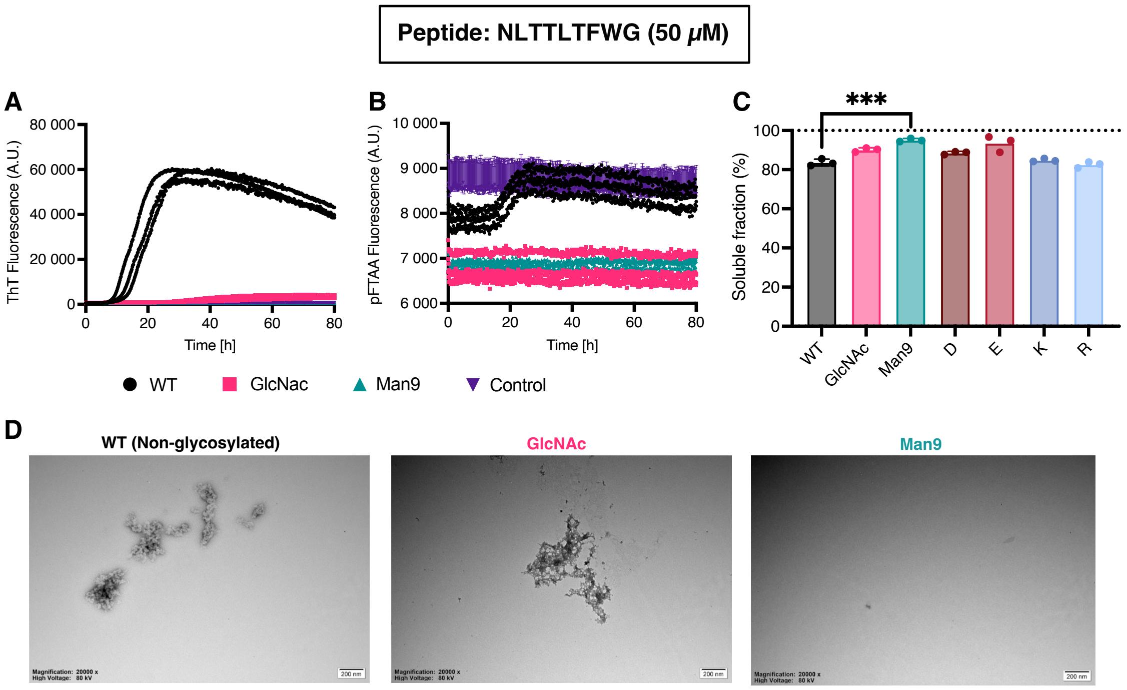

**Supplementary figure 7. A, B)** ThT binding (A) and pFTAA binding (B) kinetics. Fluorescence over time is shown for three independent repeats. Vehicle control fluorescence is shown in purple. **C)** Percentage of the concentration of peptide in the soluble fraction after ultracentrifugation (n=3). Unpaired t-test was used to assess significance of wt against Man9. **D)** TEM images after 7 days of incubation.

#### Supplementary figure 8

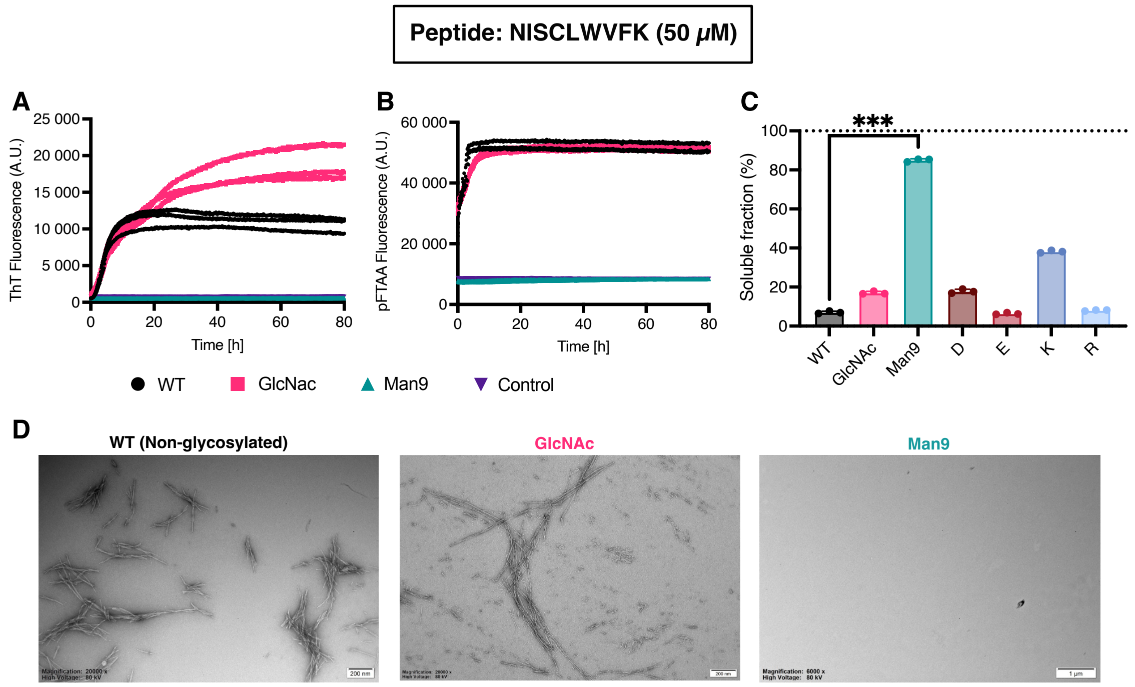

**Supplementary figure 8. A, B)** ThT binding (A) and pFTAA binding (B) kinetics. Fluorescence over time is shown for three independent repeats. Vehicle control fluorescence is shown in purple. **C)** Percentage of the concentration of peptide in the soluble fraction after ultracentrifugation (n=3). Unpaired t-test was used to assess significance of wt against Man9. **D)** TEM images after 7 days of incubation.

#### Supplementary figure 9

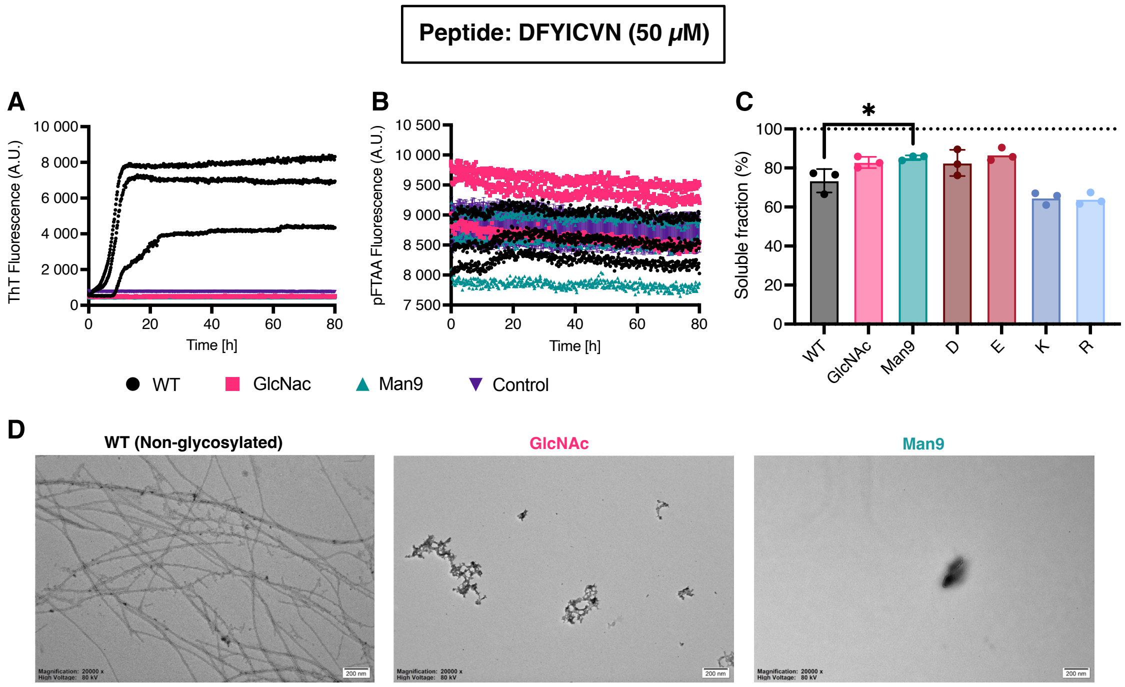

**Supplementary figure 9. A, B)** ThT binding (A) and pFTAA binding (B) kinetics. Fluorescence over time is shown for three independent repeats. Vehicle control fluorescence is shown in purple. **C)** Percentage of the concentration of peptide in the soluble fraction after ultracentrifugation (n=3). Unpaired t-test was used to assess significance of wt against Man9. **D)** TEM images after 7 days of incubation.

### Supplementary figure 10

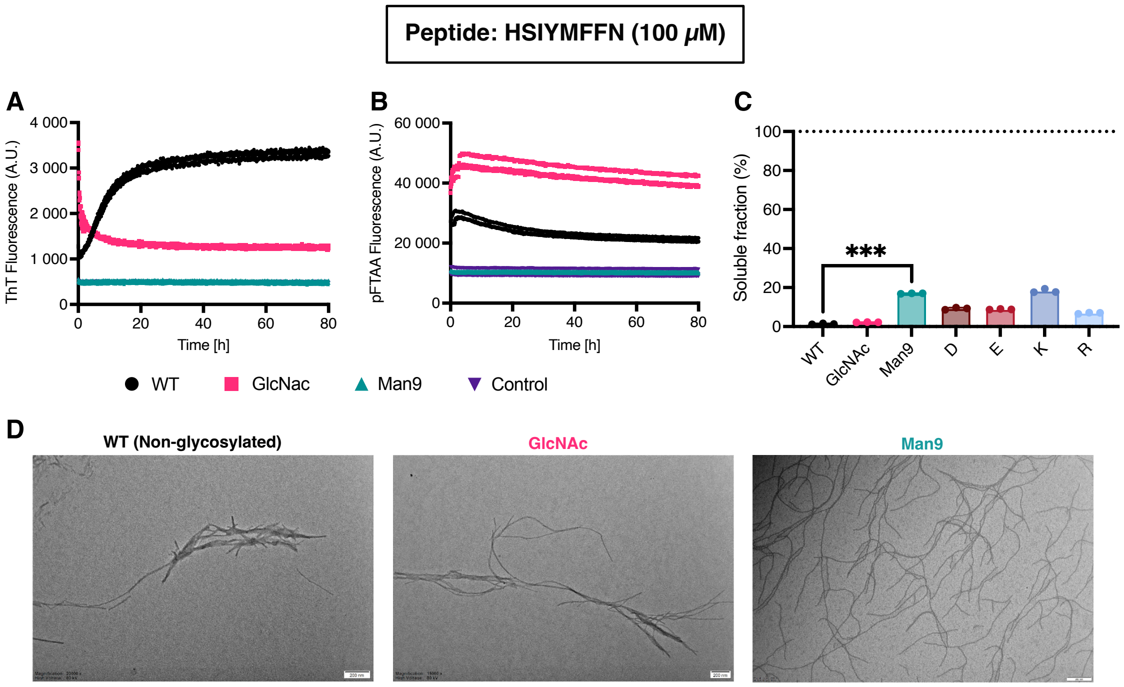

**Supplementary figure 10. A, B)** ThT binding (A) and pFTAA binding (B) kinetics. Fluorescence over time is shown for three independent repeats. Vehicle control fluorescence is shown in purple. **C)** Percentage of the concentration of peptide in the soluble fraction after ultracentrifugation (n=3). Unpaired t-test was used to assess significance of wt against Man9. **D)** TEM images after 7 days of incubation.

#### Supplementary figure 11

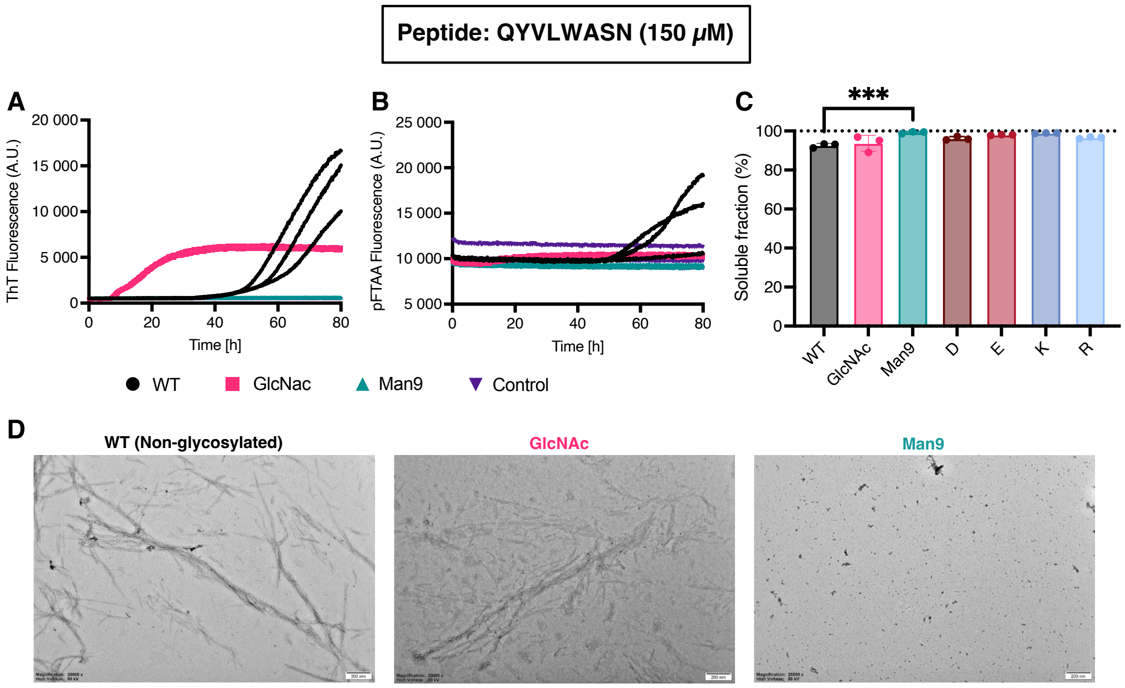

**Supplementary figure 11. A, B)** ThT binding (A) and pFTAA binding (B) kinetics. Fluorescence over time is shown for three independent repeats. Vehicle control fluorescence is shown in purple. **C)** Percentage of the concentration of peptide in the soluble fraction after ultracentrifugation (n=3). Unpaired t-test was used to assess significance of wt against Man9. **D)** TEM images after 7 days of incubation.

#### Supplementary figure 12

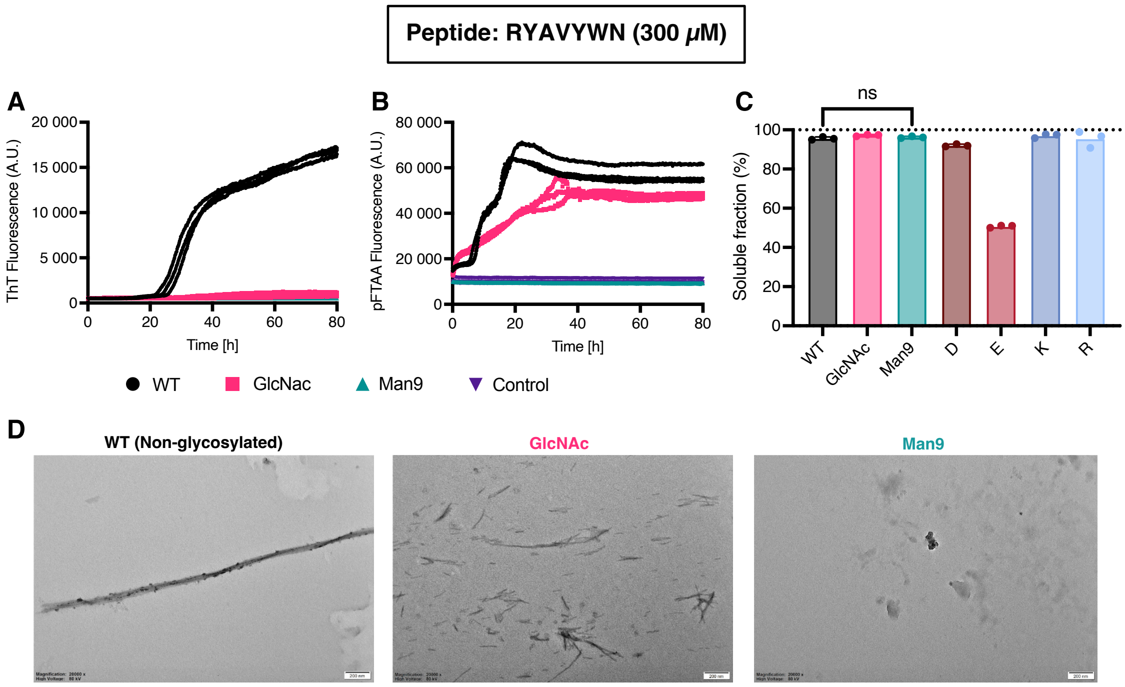

**Supplementary figure 12. A, B)** ThT binding (A) and pFTAA binding (B) kinetics. Fluorescence over time is shown for three independent repeats. Vehicle control fluorescence is shown in purple. **C)** Percentage of the concentration of peptide in the soluble fraction after ultracentrifugation (n=3). Unpaired t-test was used to assess significance of wt against Man9. **D)** TEM images after 7 days of incubation.

### Supplementary figure 13

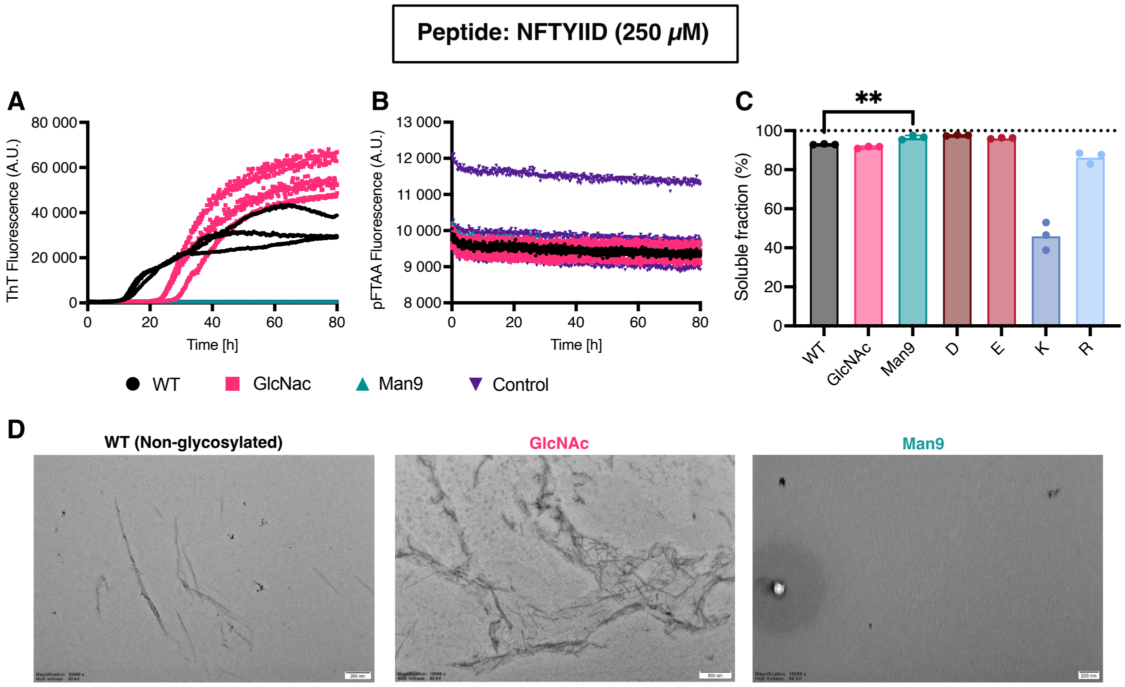

**Supplementary figure 13. A, B)** ThT binding (A) and pFTAA binding (B) kinetics. Fluorescence over time is shown for three independent repeats. Vehicle control fluorescence is shown in purple. **C)** Percentage of the concentration of peptide in the soluble fraction after ultracentrifugation (n=3). Unpaired t-test was used to assess significance of wt against Man9. **D)** TEM images after 7 days of incubation.

#### Supplementary figure 14

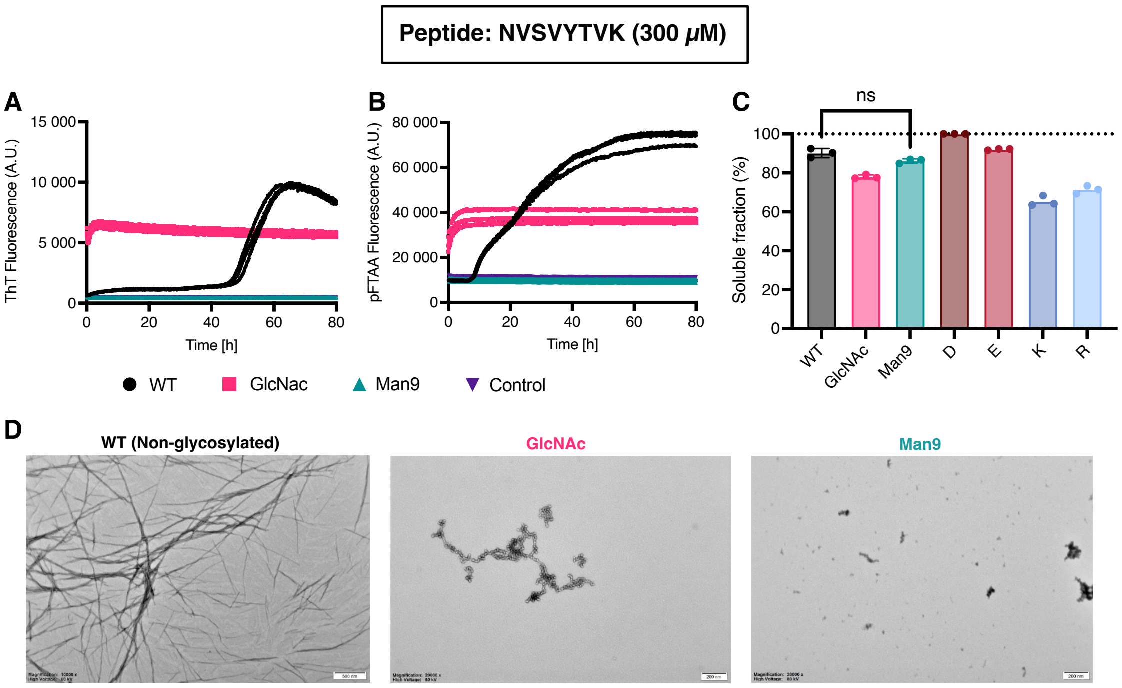

**Supplementary figure 14. A, B)** ThT binding (A) and pFTAA binding (B) kinetics. Fluorescence over time is shown for three independent repeats. Vehicle control fluorescence is shown in purple. **C)** Percentage of the concentration of peptide in the soluble fraction after ultracentrifugation (n=3). T-test was used to assess significance of wt against Man9. **D)** TEM images after 7 days of incubation.

#### Supplementary figure 15

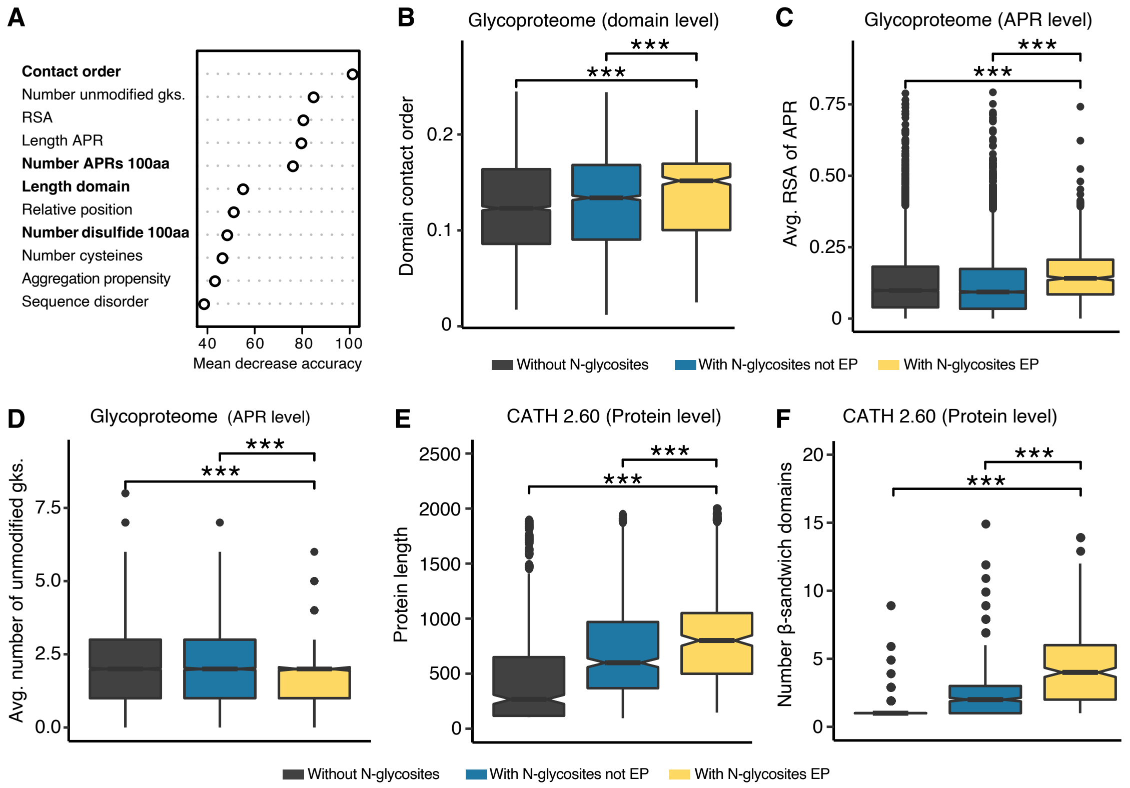

**Supplementary figure 15. A)** Variable importance plot for the model built using random oversampling (ROSE). Mean accuracy indicates the performance of the model after removing a specific variable. Higher values indicate more importance of that variable in predicting protected vs unprotected APRs. Domain-specific variables are highlighted in bold, while APR-specific variables are unhighlighted. **B-D)** Boxplot showing the relative contact order of domains (A), the average relative solvent accessibility of APRs in domains (B) or the average number of unmodified gatekeeper residues of APRs in domains (C) with N-glycosites in EPs (yellow), with N-glycosites not EPs (blue) and without N-glycosylated sites (black). Unpaired Wilcoxon test was used to assess significance among groups with Bonferroni correction for multiple comparisons. **E)** Boxplot showing the length of proteins with at least one CATH 2.60 domain with N-glycosites in EPs (yellow), with N-glycosites not EPs (blue) and without N-glycosylated sites (black). For visualization purposes, observations >2000 are not displayed in the plot. Unpaired Wilcoxon test was used to assess significance among groups with Bonferroni correction for multiple comparisons. **F)** Boxplot showing the number of b-sandwich domains in proteins with at least one CATH 2.60 domain with N-glycosites in EPs (yellow), with N-glycosites not EPs (blue) and without N-glycosylated sites (black). For visualization purposes, observations >15 are not displayed in the plot. Unpaired Wilcoxon test was used to assess significance among groups with Bonferroni correction for multiple comparisons.

#### Supplementary figure 16

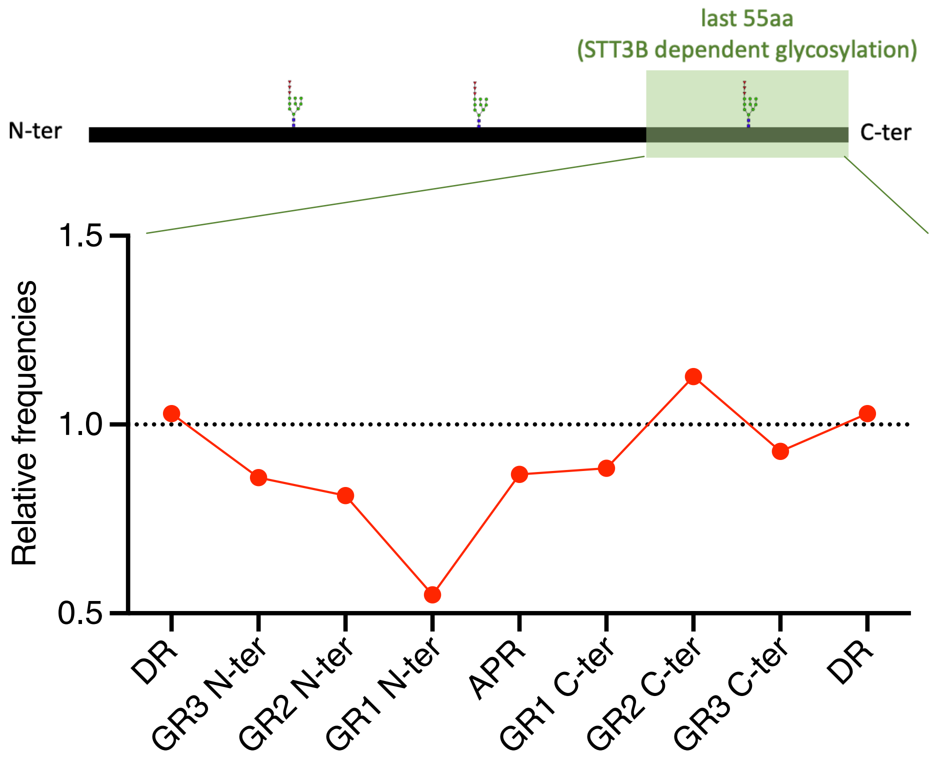

**Supplementary figure 16.** Relative frequencies of glycosylated sequons in the last 55 C-terminal residues of proteins for each region. C-terminal sites located in the last 55 residues of a protein are prone to be skipped by the translocation channel-associated STT3A isoform as they move very rapidly. Hence, extreme C-terminal glycosites are modified by an STT3B-dependent posttranslational mechanism. Glycosylated sites in EPs positions (GR2 N-ter, GR1 N-ter and APR) are significantly underrepresented ($p$ < 0.05 by Fisher exact test with FDR correction), suggesting a mechanism independent from STT3B.
